# Partial microglial depletion attenuates Purkinje cell degeneration and reshapes both glial and inflammatory responses in a mouse model of selective neurodegeneration

**DOI:** 10.64898/2026.11.14.766652

**Authors:** David Pérez-Boyero, Jimena Pérez-Arévalo, Jorge Valero, Eduardo Weruaga, David Díaz

## Abstract

**Background:** The microglial population contributes to the inflammatory environment during neurodegeneration. Its role in neuronal loss remains hard to distinguish from its response to injury. We investigated how early pharmacological reduction of this population affects Purkinje cell preservation, behaviour and the cerebellar glial response.

**Methods:** We administered PLX5622 to Purkinje cell degeneration (PCD) and wild-type (WT) mice from postnatal day (P) 15 to P24 and compared them with their respective untreated controls. We assessed motor coordination, spontaneous behaviours and novel-object recognition at P17, P21 and P25. At P25, we quantified regional Purkinje cell preservation, glial immunoreactivity and microglial abundance and morphology. We assessed selected transcripts in whole-cerebellum samples by reverse transcription quantitative polymerase chain reaction (RT–qPCR).

**Results:** At P25, treated PCD mice showed better motor coordination, improvements in selected spontaneous behaviours and greater novel-object discrimination. Greater Purkinje cell preservation coincided with lower glial fibrillary acidic protein (GFAP) immunoreactivity in lobes IV–V and VIII. Microglial density decreased in all four PCD regions analysed. The territory encompassed by the processes of residual microglia expanded in both genotypes. In PCD mice, CD68-positive tissue area decreased in lobes I–II and IV–V, and the proportion of Iba1-positive area occupied by CD68 decreased in IV–V. CD45, CD16/32 and CD206 immunoreactivity decreased alongside selective reductions in immune, inflammatory and phagocytosis-associated transcripts. Treated WT mice showed early behavioural changes, including reduced novel-object discrimination.

**Conclusions:** Early PLX5622 treatment can modify the outcome of genetically driven cerebellar degeneration in PCD mice. The benefits achieved with partial microglial depletion provide a preclinical rationale for testing how the timing and extent of glial modulation influence neuronal protection during active degeneration.

## Background

Neurodegenerative diseases are characterised by the progressive and selective loss of vulnerable neuronal populations, leading to motor, cognitive and behavioural decline [1, 2, 3]. Although their aetiology is complex and multifactorial, neuroinflammation can actively contribute to the progression of neuronal damage [2, 4, 5]. The microglial population plays a central role in this environment. Under physiological conditions, it contributes to central nervous system homeostasis through parenchymal surveillance, debris clearance, synaptic remodelling and communication with neurons and astrocytes [6, 7, 8, 9]. Persistent dysregulation of these functions can promote neurodegeneration through inflammatory signalling, oxidative stress, aberrant phagocytosis and disruption of glia–neuron communication [10, 11, 12, 13, 14].

Responses of microglia encompass diverse, dynamic and context-dependent functional states whose complexity extends beyond binary resting/activated or M1/M2 classifications [8, 15, 16, 17]. In neurodegeneration, inflammatory, phagocytic, reparative and disease-associated features may coexist [2]. Understanding how these responses interact with neuronal vulnerability and modify the neuroinflammatory environment during disease progression is therefore important [18]. Experimental models with defined patterns of neuronal loss allow these processes to be examined together and related to functional alterations.

The cerebellum has an ordered cellular architecture, well-defined circuits and a characteristic regional organisation [19]. Beyond motor coordination, it contributes to cognitive, emotional and adaptive processes [20, 21]. Cerebellar microglia have regionally specialised phenotypic and functional properties [22, 23, 24, 25] and interact dynamically with Purkinje cells, influencing neuronal excitability and cerebellar circuit function [23, 26, 27]. These features provide a useful experimental context for studying relationships between neuronal vulnerability, glial responses and behavioural alterations.

First described in 1976 [28], the Purkinje cell degeneration (PCD) mouse is a model of cerebellar neurodegeneration caused by a loss-of-function mutation in *Agtpbp1*, also known as *Ccp1/Nna1*, which encodes cytosolic carboxypeptidase 1 (CCP1) [29, 30, 31]. Biallelic pathogenic variants in its human orthologue, *AGTPBP1*, cause childhood-onset neurodegeneration with cerebellar atrophy (CONDCA), supporting the translational relevance of the model [32, 33]. PCD mice develop selective Purkinje cell degeneration during the early postnatal period [28, 34], with regionally heterogeneous progression and relative resistance of lobe X [35, 36, 37]. This process is accompanied by ataxia and motor impairment [38], cerebellar glial alterations [39], selective leucocyte recruitment and peripheral immune changes [40]. Its defined progression and regional differences in vulnerability allow investigation of how glial responses interact with a primary process of neuronal degeneration.

Our previous work showed that CCP1-deficient microglia exhibit intrinsic and environment-dependent alterations in phagocytosis, motility and proliferation, together with an atypical inflammatory profile enriched in markers traditionally associated with anti-inflammatory and reparative responses in degenerating regions [41]. Microgliosis, astrogliosis and increased expression of glial and inflammatory markers have also been described in the PCD cerebellum [39, 42]. Differences between the cerebellum and olfactory bulb, another region affected in this model, indicate that glial responses depend on region and disease stage [39, 41]. In other experimental models, specific microglial signals can induce astroglial responses that contribute to neuronal damage [43, 44]. These findings justify examining how reducing the microglial population modifies the broader glial and inflammatory environment during cerebellar degeneration.

Pharmacological manipulation of the microglial population allows investigation of its contribution to neurodegenerative processes [45, 46]. PLX5622, a colony-stimulating factor 1 receptor (CSF1R) inhibitor, is used to substantially reduce this population and examine its involvement in disease progression [47, 48, 49]. CSF1R inhibition can produce marked but incomplete microglial depletion [47, 50, 51]. Cells remaining after treatment may exhibit distinct morphological and molecular features [50, 51], supporting investigation of residual microglia and their relationship with changes in the glial environment. Systemic CSF1R inhibition may also affect peripheral myeloid and immune compartments, which should be considered when interpreting the effects of PLX5622 [52, 53].

Our previous study linked pharmacological reduction of the microglial population to increased Purkinje cell survival and improved motor performance in PCD mice, with the motor benefit evaluated at postnatal day (P) 25 [41]. The present study extends this evidence by examining multiple behavioural domains over the degenerative period and characterising treatment-associated glial responses in greater depth. We ask how the functional effects of PLX5622 evolve over time and which regional, cellular and molecular changes accompany neuronal preservation.

In the present study, we combine an expanded behavioural battery, a larger rotarod cohort and longitudinal assessments at P17, P21 and P25. In addition to motor coordination, we assess spontaneous behaviours—grooming, rearing, walking and resting—and novel-object recognition. At P25, this functional assessment is integrated with regional analysis of Purkinje cell preservation and detailed glial characterisation. The glial analysis addresses residual microglial abundance and morphology, together with markers associated with glial and immune reactivity and with proinflammatory and anti-inflammatory/reparative responses. We also evaluate lysosomal immunoreactivity associated with phagocytic processes at the tissue level and within the microglial compartment, as well as astroglial reactivity. These analyses are complemented by targeted analysis of glial and inflammatory gene expression in the whole cerebellum. This approach allows joint examination of behavioural trajectories and the neuroimmune environment associated with neuronal preservation during active cerebellar degeneration.

## Materials and methods

### Experimental animals and genotyping

Wild-type (WT) and Purkinje cell degeneration (PCD) mutant mice (*Mus musculus*) on a C57BL/DBA genetic background were used in this study. PCD mice were homozygous for the *pcd1J* allele of *Agtpbp1* (also known as *Ccp1/Nna1*) [42]. Animals were housed under standard laboratory conditions, with controlled temperature and humidity, a 12 h light/dark cycle, and ad libitum access to food and water.

All experimental procedures were conducted in accordance with current European legislation governing the use of laboratory animals, including Directive 2010/63/EU and Recommendation 2007/526/EC, as well as Spanish regulations, including Law 32/2007 and Royal Decree 53/2013. The experimental protocol was approved by the Bioethics Committee of the University of Salamanca, Spain, under reference numbers #00613 and #1258.

The colony was maintained by breeding heterozygous animals. Offspring were genotyped by polymerase chain reaction following previously established protocols [54, 55]. Within each genotype, animals were randomly allocated to the treatment and control groups. Predefined exclusion criteria included unsuccessful genotyping, technical failure during behavioural acquisition, poor tissue preservation, incomplete cerebellar lobe anatomy preventing reliable quantification, or insufficient RNA quality for reverse transcription quantitative polymerase chain reaction (RT–qPCR). Exclusion criteria were applied before unblinding whenever applicable.

### Experimental design

Four experimental groups were analysed: untreated WT, PLX5622-treated WT, untreated PCD and PLX5622-treated PCD mice. The accelerating rotarod test was performed using *n* = 8 animals per group, whereas spontaneous behaviour and novel object recognition were assessed using *n* = 4 animals per group. After behavioural assessment, half of the animals in each group were assigned to histological/immunohistochemical analyses and the remaining half to targeted gene expression analyses, with *n* = 4 animals per group assigned to each experimental subset. For all analyses, the individual animal was considered the experimental unit.

PLX5622 treatment was administered from postnatal day (P) 15 to P24, covering the period of maximal Purkinje cell vulnerability in the PCD model. Behavioural testing was performed longitudinally at P17, P21 and P25 to evaluate motor coordination, spontaneous behaviour and recognition performance. The same animals underwent behavioural assessment at all three time points within each behavioural experiment. After completion of behavioural testing and treatment administration, animals were euthanised at P25 and processed for immunohistochemical or targeted gene expression analyses. All immunohistochemical and targeted gene expression analyses were therefore performed at P25.

PLX5622-treated animals were compared with age- and genotype-matched untreated controls. Immunohistochemical analyses were performed in the cerebellar vermis, focusing on selected lobes according to the specific endpoint analysed and the regional vulnerability pattern of the PCD model at this age [35, 36, 37]. Lobes I–II, IV–V, VIII and X were included when appropriate to assess region-dependent differences in Purkinje cell degeneration and glial responses. The exact lobes analysed for each quantification are specified in the corresponding figure legends.

Targeted gene expression analyses were performed using whole-cerebellum samples to evaluate the overall impact of PLX5622 treatment on selected transcripts associated with microglial identity, inflammatory signalling, phagocytosis, motility and chemotaxis.

Behavioural scoring, histological and immunohistochemical quantification, image analysis and statistical analyses were performed under blinded experimental conditions whenever possible. Group identities were revealed only after completion of the corresponding analyses.

### Pharmacological microglial depletion

PLX5622, a selective colony-stimulating factor 1 receptor (CSF1R) inhibitor (MedChemExpress, Monmouth Junction, NJ, USA), was administered intraperitoneally using corn oil as the vehicle because of its poor solubility [41]. Injection was necessary because independent consumption of medicated chow could not be ensured before weaning. Previous studies administered 50 mg/kg PLX5622 intraperitoneally to neonatal rats and mice [56, 57, 58, 59]. Four 50 mg/kg doses were given 72 h apart at P15, P18, P21 and P24 [41]. Starting at P15 limited exposure during earlier cerebellar organisation and circuit formation. The regimen began before overt Purkinje cell loss and covered the period of maximal vulnerability [41]. Pilot experiments guided the choice of starting age and injection interval to reduce microglial abundance while preserving animal welfare [41]. Earlier initiation or more frequent injections caused weight loss, lethargy and increased mortality, so these schedules were excluded. The selected schedule was well tolerated in those pilots, without distress, overt behavioural abnormalities or weight loss [41]. It produced substantial but incomplete microglial depletion, consistent with reports that short-term CSF1R inhibition leaves a residual population and that near-complete elimination generally requires longer treatment [47, 50, 51].

### Behavioural assessment

Behavioural assessments used protocols adapted from previous work in our laboratory [38, 41, 60, 61]. All behavioural assessments were performed longitudinally at P17, P21 and P25.

Motor coordination and balance were assessed using an accelerating RotaRod LE8200 apparatus (Panlab, Cornellà de Llobregat, Barcelona, Spain; Figure 1A). The same animals were tested longitudinally at P17, P21 and P25. Animals were placed on a rotating rod with a diameter of 3.5 cm, which accelerated progressively from 4 to 40 rpm over a 10 min period. Latency to fall was automatically recorded. Each animal performed seven trials per session, separated by 10 min rest intervals, and the mean latency to fall was calculated for each analysed age.

**Figure 1.**
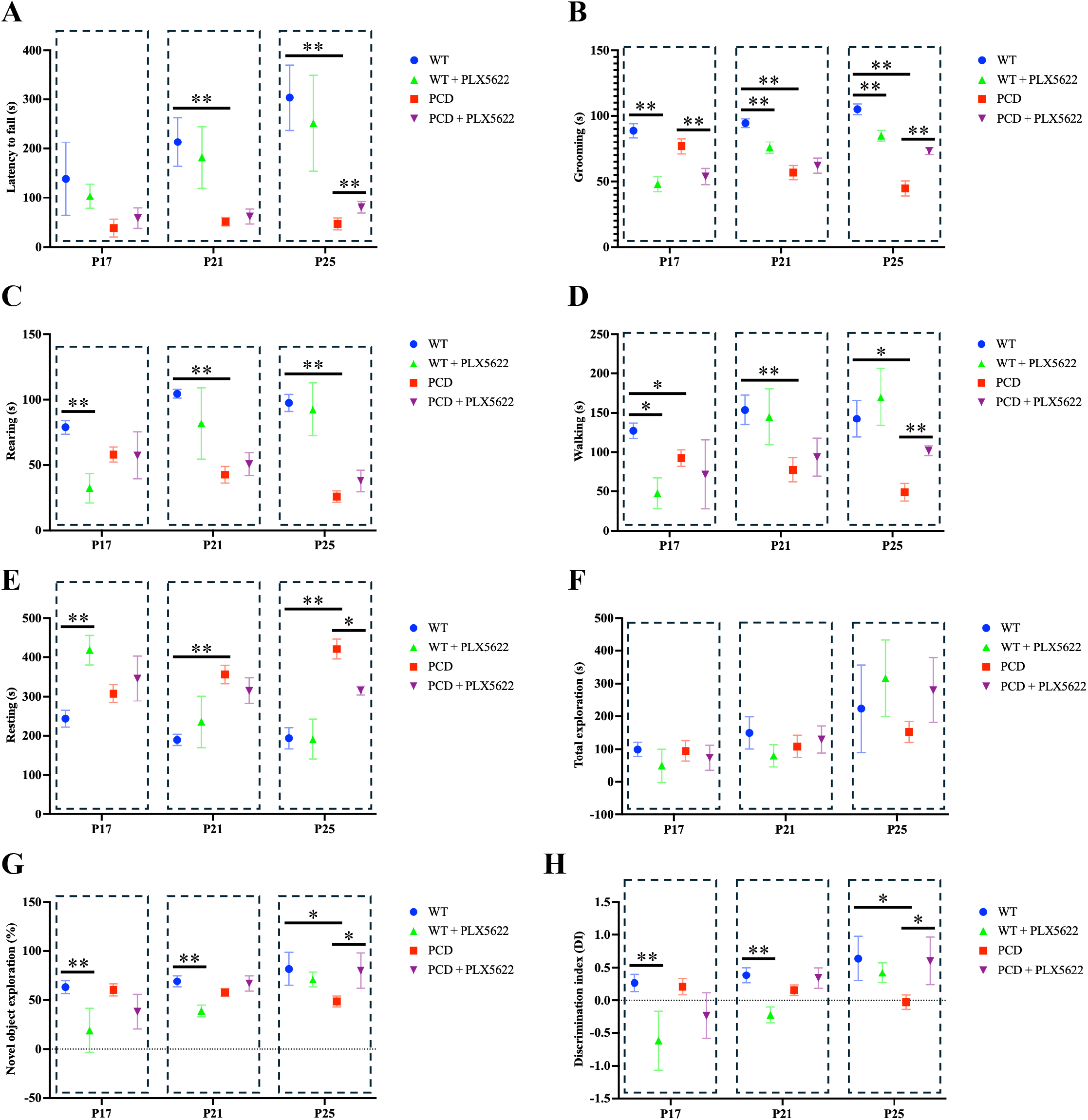
Effects of PLX5622 on motor performance, spontaneous behaviour and novel-object recognition. Wild-type (WT) and Purkinje cell degeneration (PCD) mice, with or without PLX5622 treatment, were assessed longitudinally at postnatal days (P) 17, 21 and 25. (A) Motor coordination, measured as latency to fall (s) from an accelerating rotarod. (B–E) Time spent (s) grooming (B), rearing (C), walking (D) and resting (E) during spontaneous behavioural assessment. (F–H) Novel-object recognition, expressed as total exploration time (s) for both objects (F), percentage of object exploration time devoted to the novel object (G), and discrimination index (H). Symbols and error bars represent group means and SD, respectively; *n* = 8 animals per group in A and *n* = 4 per group in B–H. Each animal was assessed at all three ages. Data were analysed using two-way repeated-measures ANOVA, with experimental group and age as factors. Bonferroni correction was applied to the three prespecified comparisons (WT versus PCD, WT versus WT + PLX5622, and PCD versus PCD + PLX5622) within each age and behavioural endpoint. Brackets identify the indicated between-group comparisons; \**p* < 0.05; \*\**p* < 0.01.

General spontaneous behaviour was evaluated in standard Sealsafe® cages (Tecniplast S.p.A., Buguggiate, Italy) containing a 1.5-cm layer of bedding material. After a 10 min habituation period, animals were video recorded for an additional 10 min. Recordings were subsequently analysed to quantify the time spent grooming, rearing, walking and resting (Figure 1B–E).

Recognition performance was assessed using the novel object recognition test (Figure 1F–H). During the familiarisation phase, animals were allowed to explore two identical objects for 10 min. After a 10 min retention interval, one object remained familiar, whereas the other was replaced by a novel object for the test phase. Animals were then allowed to explore freely for 10 min. Exploration was defined as the animal being oriented towards the object at a distance of less than 2 cm. The time spent exploring the novel object (TN) and familiar object (TF) was recorded. Total object exploration time was defined as TN + TF. The percentage of total object exploration time spent exploring the novel object was calculated as:

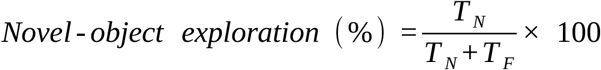

The discrimination index (DI) was calculated as:

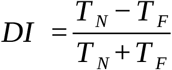

Positive DI values indicated greater exploration of the novel object, negative values indicated greater exploration of the familiar object, and zero indicated equal exploration. These descriptive values did not by themselves establish statistically significant discrimination from chance. Total exploration time was analysed to determine whether differences in recognition performance could be influenced by altered object-directed exploratory activity.

### Tissue processing and immunohistochemistry

Animals assigned to histological analyses were deeply anaesthetised and transcardially perfused using a peristaltic pump (Gilson, Middleton, WI, USA). The vascular system was first rinsed with 0.9% isotonic saline solution and subsequently perfused for 15 min with glutaraldehyde-free Somogyi’s fixative containing 4% (w/v) paraformaldehyde (VWR Chemicals, Solon, OH, USA) and 0.2% (v/v) saturated picric acid (Probus S.A., Badalona, Spain) in 0.1 M phosphate buffer (PB), pH 7.4.

Cerebella were frozen and sectioned parasagittally at 30 μm using a Jung SM 2000 sliding freezing microtome (Leica Instruments, Nussloch, Germany). Sections corresponding to the cerebellar vermis and containing complete cerebellar lobe anatomy were selected for analysis.

Free-floating sections were incubated for 24 h at 4 °C under continuous gentle agitation in phosphate-buffered saline (PBS) containing 0.2% (v/v) Triton X-100 (Probus S.A.), 5% (v/v) normal donkey serum (Vector Laboratories, Burlingame, CA, USA) and the corresponding primary antibodies. After washing in PBS, sections were incubated for 1 h at room temperature with the appropriate secondary antibodies. 4′,6-Diamidino-2-phenylindole (DAPI; Sigma-Aldrich, St. Louis, MO, USA) was added 10 min before the end of the secondary antibody incubation at a dilution of 1:10,000. Finally, sections were mounted onto gelatine-coated slides and coverslipped using an antifade mounting medium.

Negative controls were processed in parallel by omitting primary antibodies, and no specific signal was detected. Primary and secondary antibodies, including host species, suppliers, catalogue numbers and working dilutions, are listed in Table 1.

**Table 1.** Primary and secondary antibodies used for cerebellar immunofluorescence.

| **Primary antibody** | **Host** | **Source / catalogue no.** | **Dilution** | **Secondary antibody** | **Source / catalogue no.** | **Dilution** |
|---|---|---|---|---|---|---|
| Calbindin D-28k | Mouse | Swant (Kehrsatz, Switzerland); CB300 | 1:1000 | Cy3 donkey anti-mouse IgG (H+L) | Jackson ImmunoResearch (West Grove, PA, USA); 715-165-151 | 1:500 |
| CD16/32 | Rat | BD Biosciences (San Jose, CA, USA); 553142 | 1:200 | Cy3 donkey anti-rat IgG (H+L) | Jackson ImmunoResearch; 712-165-153 | 1:500 |
| CD206 | Goat | R&D Systems (Minneapolis, MN, USA); AF2535 | 1:200 | Cy3 donkey anti-goat IgG (H+L) | Jackson ImmunoResearch; 705-165-147 | 1:500 |
| CD45 | Rat | Bio-Rad (Hercules, CA, USA); MCA1388 | 1:1000 | Cy3 donkey anti-rat IgG (H+L) | Jackson ImmunoResearch; 712-165-153 | 1:500 |
| CD68 | Rat | Thermo Fisher Scientific (Waltham, MA, USA); 14-0681-82 | 1:500 | Cy5 donkey anti-rat IgG (H+L) | Jackson ImmunoResearch; 712-175-153 | 1:500 |
| Iba1 | Rabbit | FUJIFILM Wako (Osaka, Japan); 019-19741 | 1:1000 | Cy2 or Cy3 donkey anti-rabbit IgG (H+L) | Jackson ImmunoResearch; 711-225-152 / 711-165-152 | 1:500 |
| GFAP | Mouse | Sigma-Aldrich; G3893 | 1:2000 | Cy3 donkey anti-mouse IgG (H+L) | Jackson ImmunoResearch; 715-165-151 | 1:500 |

GFAP, glial fibrillary acidic protein; Iba1, ionised calcium-binding adaptor molecule 1; H+L, heavy and light chains.
For Iba1, the secondary fluorophore depended on the antibody combination used.

Microglia were identified based on Iba1 immunoreactivity, characteristic morphology and anatomical localisation within the cerebellar parenchyma. CD68 was analysed as a marker associated with lysosomal and phagocytic activity, whereas CD45 was evaluated as a marker associated with immune and microglial reactivity. CD16/32 and CD206 were analysed as markers associated with proinflammatory and anti-inflammatory/reparative responses, respectively, without implying a binary M1/M2 classification. Astroglial reactivity was assessed by GFAP immunoreactivity.

### Image acquisition and quantitative analyses

Fluorescence images were acquired using a STELLARIS 8 confocal microscope (Leica Microsystems, Wetzlar, Germany). Acquisition settings and microscope configurations were adjusted according to the requirements of each specific analysis and were kept constant across experimental groups within the same experiment to allow direct comparison between conditions.

For morphological analyses, confocal z-stacks were acquired and processed as maximum-intensity projections before quantification.

Quantitative analyses were performed in cerebellar lobes I–II, IV–V, VIII and X when appropriate. Instead of analysing a fixed number of microscopic fields per animal, quantifications were performed across the full extent of each analysed lobe to minimise sampling bias and account for regional heterogeneity.

Microglial density, Purkinje cell density, immunoreactive area, integrated density and co-localisation analyses were performed using ImageJ (National Institutes of Health, Bethesda, MD, USA) [62] with the Fiji distribution [63]. Purkinje cells were identified by calbindin D-28k immunoreactivity and quantified along the Purkinje cell layer to obtain linear cell density (cells/mm). Calbindin D-28k-positive area fraction was also quantified as a measure of Purkinje cell immunoreactivity.

All identified microglial cells were counted throughout each complete cerebellar section analysed, with counts for the predefined lobes retained separately. Microglial abundance was quantified using microglial density (cells/mm3) and the percentage of analysed area occupied by Iba1-positive immunoreactivity. GFAP, CD45, CD16/32 and CD206 immunoreactivity were quantified as the percentage of the analysed area occupied by positive staining. CD68 immunoreactivity was quantified both as the percentage of the analysed tissue area within each lobe occupied by CD68-positive staining and as the percentage of Iba1-positive microglial area occupied by CD68 immunoreactivity.

For regional measurements of Purkinje cell preservation, microglial abundance, CD68 and GFAP, each predefined lobe was analysed separately. For CD45, CD16/32 and CD206, the percentages of positively stained area in lobes I–II, IV–V, VIII and X were averaged within each animal to obtain a single mean percentage per marker. Individual Purkinje cell and microglial morphological measurements were combined across the sampled lobes within each animal before group comparisons.

The morphology of surviving Purkinje cells was analysed by manually segmenting individual neurons and quantifying soma area, dendritic tree area, dendritic territory area and mean calbindin D-28k fluorescence intensity using Fiji/ImageJ. Dendritic tree area was defined as the area occupied by the labelled dendritic processes in the analysed projection. Dendritic territory area was defined as the total tissue area encompassed by the dendritic arbour, including the spaces between its processes. Quantitative single-cell analyses were restricted to untreated and PLX5622-treated PCD mice because extensive overlap of dendritic labelling in WT and WT + PLX5622 mice prevented reliable delineation of individual dendritic trees.

Microglial morphology was analysed in Iba1-positive cells using Fiji/ImageJ to quantify cell area, soma area, the territory encompassed by the processes (tracking area) and Iba1 integrated density. Tracking area describes the total area enclosed by the processes, including the spaces between them.

Thresholding and quantification parameters were kept constant within each analysis. All image analyses were performed under blinded experimental conditions. For each animal, values obtained from the analysed sections, cells and lobes were averaged as appropriate before statistical analysis, ensuring that the animal, rather than individual images or cells, was used as the experimental unit.

### RNA extraction and RT–qPCR

Animals assigned to targeted gene expression analyses were deeply anaesthetised and transcardially perfused with 0.9% isotonic saline solution. Whole cerebella were rapidly dissected using RNase-free material and immediately processed.

Tissue samples were homogenised using an Ultra-Turrax homogeniser (IKA, Staufen, Germany). Total RNA was extracted and purified using the PureLink™ RNA Mini Kit together with the PureLink™ DNase Set (Thermo Fisher Scientific), according to the manufacturer’s instructions.

RNA was reverse transcribed into cDNA using the High-Capacity cDNA Reverse Transcription Kit (Applied Biosystems, Foster City, CA, USA). RT–qPCR analyses were performed using a QuantStudio 7 Flex Real-Time PCR System (Applied Biosystems), following previously described protocols [42, 64, 65, 66]. Relative gene expression levels were normalised to *Gapdh* and calculated using the 2*^−ΔΔCt^* method [67].

Four biological replicates, corresponding to four individual animals, were analysed per experimental group. Genes were organised into functional groups comprising: (1) microglia-associated markers, including *Iba1*, *Cx3cr1* and *Tmem119*; (2) phagocytosis-associated genes, including *Trem2*, *P2ry6* and *Cd68*; (3) motility- and chemotaxis-associated genes, including *Ccl2* and *Ccr2*; and (4) inflammatory and regulatory genes, including *Tlr2*, *Tlr4*, *Nos2* (*iNOS*), *Tnf* (*Tnf-α*), *Il1b* (*Il-1β*), *Arg1* and *Cd206*.

For the main analysis, gene expression was expressed as fold change relative to the untreated WT group.

Primer sequences are listed in Table 2.

**Table 2.**
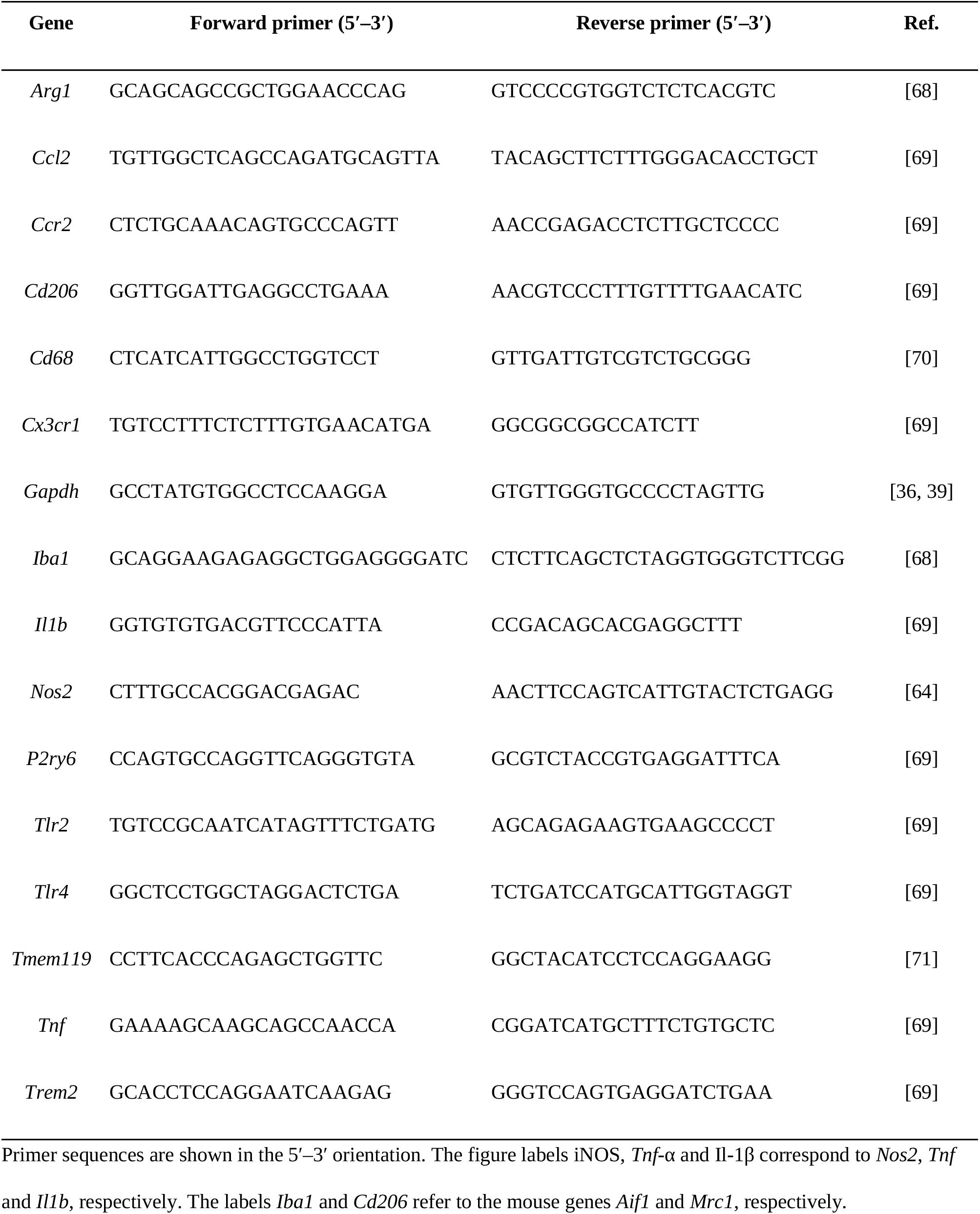
Primer sequences used for targeted RT–qPCR analysis of cerebellar tissue.

### Statistical analyses

Data are presented as mean ± standard deviation (SD), unless otherwise specified. The animal was the experimental unit, with microscopy measurements averaged per animal and, where applicable, per lobe. Analyses were performed using GraphPad Prism 10 (GraphPad Software, Boston, MA, USA) and IBM SPSS Statistics 32 (IBM Corp., Armonk, NY, USA); significance was set at p < 0.05.

RT–qPCR data were analysed by analysis of variance (ANOVA) using untransformed fold changes. The corresponding plots display geometric means with geometric SD on logarithmic axes; heat maps display geometric mean fold changes. For Supplementary Figure S1 (Additional file 1), the same expression values were additionally referenced to the geometric mean of the corresponding untreated genotype.

Longitudinal behavioural outcomes were analysed by two-way repeated-measures ANOVA, with experimental group as the between-subject factor and age (P17, P21 and P25) as the within-subject factor. Histological, immunohistochemical and RT–qPCR endpoints were analysed by ordinary two-way ANOVA, with genotype and treatment as between-subject factors. Both models included main effects and their interaction. Regional endpoints and individual genes were analysed separately.

Bonferroni correction was applied to three prespecified comparisons—WT vs PCD, WT vs WT + PLX5622, and PCD vs PCD + PLX5622—as one family per behavioural outcome at each age, or per histological/immunohistochemical endpoint (within each lobe, where applicable) or gene.

Single-cell morphological parameters of surviving Purkinje cells were compared between untreated and PLX5622-treated PCD mice using a two-tailed Mann–Whitney U test on animal-level means.

## Results

### PLX5622 improves motor coordination, selected spontaneous behaviours and novel-object discrimination in PCD mice at P25

Behaviour was assessed longitudinally at P17, P21 and P25 by analysing motor coordination, spontaneous behaviour and novel-object discrimination (Figure 1). Rotarod performance showed significant effects of experimental group and age, together with a group × age interaction (all *p* < 0.001; Figure 1A). Untreated PCD mice had a shorter latency to fall than WT mice at P21 and P25 (both *p* < 0.001), whereas no significant difference was detected at P17 (*p* = 0.057). In PCD mice, PLX5622 significantly increased latency to fall at P25 (*p* < 0.001), but not at P17 (*p* = 0.533) or P21 (*p* = 1.000). Treatment did not significantly alter WT performance at any of the ages assessed.

Spontaneous-behaviour analysis revealed significant interactions between experimental group and age for time spent grooming, rearing, walking and resting (all *p* < 0.001; Figure 1B–E). At P17, untreated PCD mice walked less than WT mice (*p* = 0.026), with no significant differences in the other behaviours. At P21 and P25, PCD mice showed less grooming and rearing—both *p* < 0.001 at both ages—and less walking (*p* = 0.008 and *p* = 0.013, respectively), together with more time resting—*p* < 0.001 at both ages. In PCD mice, PLX5622 reduced grooming at P17 (*p* < 0.001), without significantly altering the other behaviours, and produced no significant differences in any of these parameters at P21. At P25, treatment increased grooming (*p* < 0.001) and walking (*p* = 0.005) and reduced resting time (*p* = 0.014), while rearing did not change significantly. In WT mice, PLX5622 reduced grooming at all three ages—P17 and P25, *p* < 0.001; P21, *p* = 0.001. At P17, it also reduced rearing (*p* < 0.001) and walking (*p* = 0.011) and increased resting (*p* = 0.006). These latter behaviours showed no significant treatment-associated differences in WT mice at P21 or P25.

In the novel-object recognition test, total exploration time was influenced by age (*p* < 0.001), but not by experimental group (*p* = 0.341), and the group × age interaction was not significant (*p* = 0.116; Figure 1F). None of the prespecified between-group comparisons showed significant differences at the ages assessed. In particular, total exploration time did not differ significantly between treated and untreated PCD mice at P25 (*p* = 0.685).

The percentage of exploration time spent with the novel object and the discrimination index were analysed as two linearly related expressions of the same recognition outcome (Figure 1G, H). At P17 and P21, untreated WT and PCD mice did not differ significantly in these measures (*p* = 1.000 and *p* = 0.161, respectively). However, PLX5622 reduced both parameters in WT mice at P17 (*p* = 0.009) and P21 (*p* = 0.004), with no significant treatment-associated differences in PCD mice at those ages. At P25, untreated PCD mice had lower values than WT mice (*p* = 0.023), whereas PLX5622-treated PCD mice showed a higher novel-object exploration percentage and discrimination index than untreated PCD mice (*p* = 0.043). No significant treatment-associated differences were detected in WT mice at this age.

Together, these results indicate that PLX5622 attenuates several behavioural alterations in PCD mice at P25. The combination of improved motor coordination, favourable changes in selected spontaneous behaviours and greater novel-object discrimination supports a functional benefit extending across different behavioural domains. Greater discrimination was also detected without significant differences in total exploration time. This benefit in PCD, together with the early alterations observed in WT mice, shows that the intervention produces a pattern of effects that should be interpreted according to experimental group and age, rather than a uniform behavioural improvement.

### PLX5622 favours regional Purkinje cell preservation in PCD mice

Calbindin D-28k labelling showed a broad, continuous distribution in the cerebellar cortex of WT and WT + PLX5622 mice. In untreated PCD mice, the signal was sparser and more fragmented in affected regions, while lobe X retained relatively preserved labelling. Treated PCD mice showed more extensive labelling in certain regions, although sparsely labelled areas remained (Figure 2A–D).

**Figure 2.**
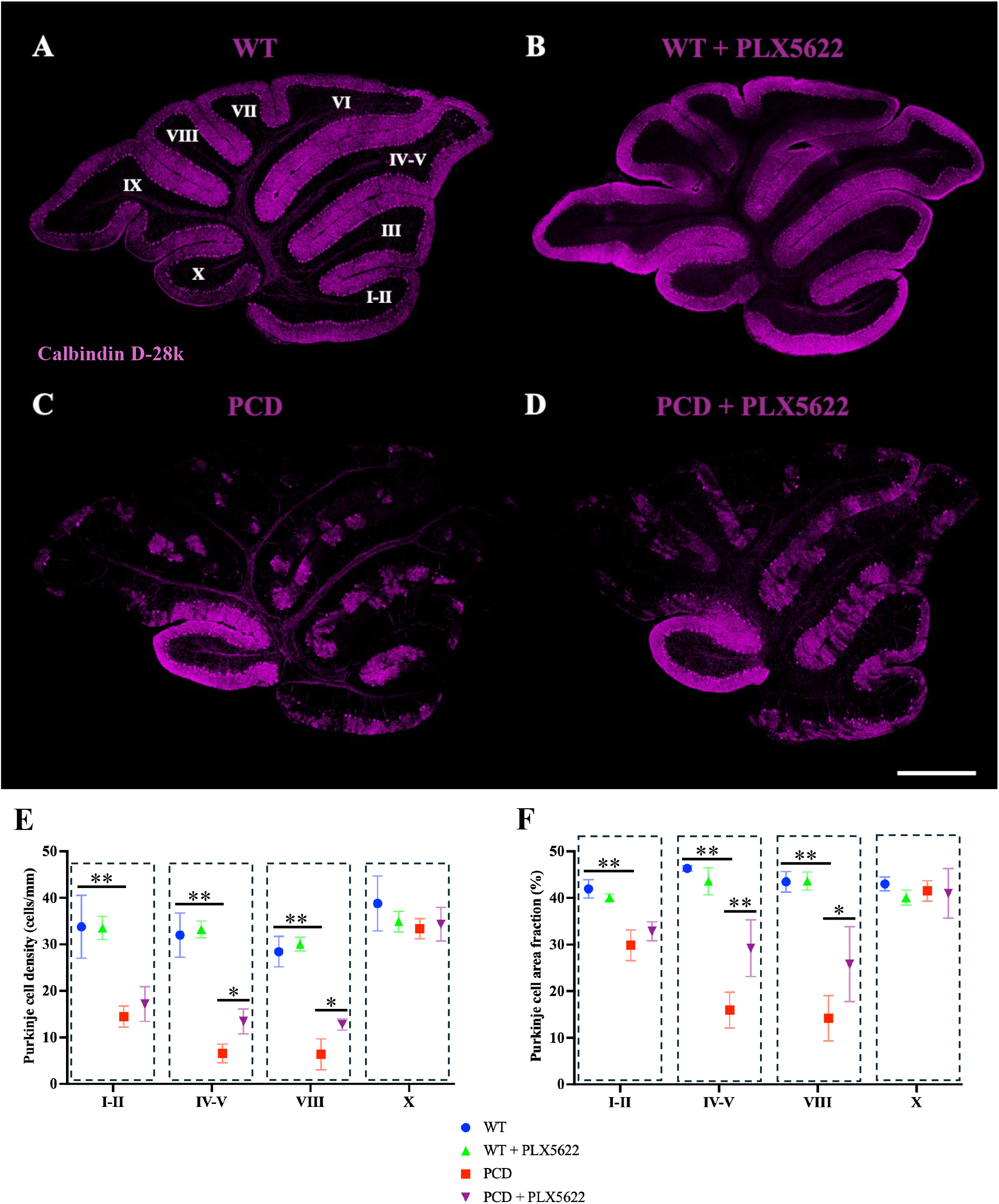
PLX5622 treatment partially preserves Purkinje cells in selected cerebellar lobes of PCD mice. (A–D) Representative confocal images of calbindin D-28k immunoreactivity (magenta) in parasagittal sections of the cerebellar vermis at P25: WT (A), WT + PLX5622 (B), PCD (C) and PCD + PLX5622 (D). (E, F) Purkinje cell linear density (cells/mm; E) and calbindin D-28k-positive area fraction (%; F) in lobes I–II, IV–V, VIII and X. Measurements were summarised within each animal for each lobe before group comparisons. Symbols and error bars represent group means and SD; *n* = 4 independent animals per group. Each lobe was analysed separately using ordinary two-way ANOVA, with genotype and treatment as factors. Bonferroni correction was applied to the three prespecified comparisons (WT versus PCD, WT versus WT + PLX5622, and PCD versus PCD + PLX5622) within each lobe and endpoint. Brackets identify the indicated between-group comparisons; \**p* < 0.05; \*\**p* < 0.01. Scale bar, 500 µm. WT, wild type; PCD, Purkinje cell degeneration.

Purkinje cell preservation was assessed at P25 by quantifying linear density and the calbindin D-28k-positive area fraction in cerebellar vermis lobes I–II, IV–V, VIII and X (Figure 2). Untreated PCD mice had a lower Purkinje cell density than WT mice in lobes I–II, IV–V and VIII (all *p* < 0.001), whereas no significant difference was detected in lobe X (*p* = 0.199). In PCD mice, PLX5622 significantly increased neuronal density in lobes IV–V (*p* = 0.022) and VIII (*p* = 0.011), with no significant differences in I–II or X. Treatment did not significantly alter Purkinje cell density in WT mice either (Figure 2E). Two-way ANOVA showed significant effects of genotype in I–II, IV–V and VIII (all *p* < 0.001) and of treatment in IV–V (*p* = 0.020) and VIII (*p* = 0.008). The genotype × treatment interaction did not reach statistical significance in any of the lobes analysed.

Calbindin D-28k-positive area showed a concordant pattern. Untreated PCD mice had a lower positive area fraction than WT mice in lobes I–II, IV–V and VIII (all *p* < 0.001), with no significant difference in X. In PCD mice, PLX5622 increased this parameter in IV–V (*p* = 0.001) and VIII (*p* = 0.017), whereas no significant changes were detected in I–II or X. No significant treatment-associated differences were observed in WT mice either (Figure 2F). The overall analysis revealed significant effects of genotype in I–II, IV–V and VIII (all *p* < 0.001) and of treatment in IV–V (*p* = 0.019) and VIII (*p* = 0.033). Genotype × treatment interactions were also detected in I–II (*p* = 0.048), IV–V (*p* = 0.001) and VIII (*p* = 0.038), but not in X.

Together, the increases in Purkinje cell density and calbindin D-28k-positive area fraction support greater preservation of this neuronal population in lobes IV–V and VIII of treated PCD mice at P25. The agreement between these parameters provides anatomical evidence of neuronal preservation accompanying the behavioural improvement at this age. Thus, the functional benefit of treatment coexists with lower Purkinje cell loss in cerebellar regions affected by degeneration.

### Greater Purkinje cell preservation is not accompanied by significant differences in the morphological parameters analysed

Qualitative examination showed that, in WT and PLX5622-treated WT mice, somata were distributed along the Purkinje cell layer and extensive, densely branched dendritic arbours overlapped widely within the molecular layer (Figure 3A–H). PCD mice showed areas with more isolated cells, whose dendritic trees varied in extent and organisation (Figure 3I–L). This heterogeneity persisted in PLX5622-treated PCD mice: cells with broad arbours coexisted with profiles displaying more restricted dendritic trees and some apparently small somata (Figure 3M–P).

**Figure 3.**
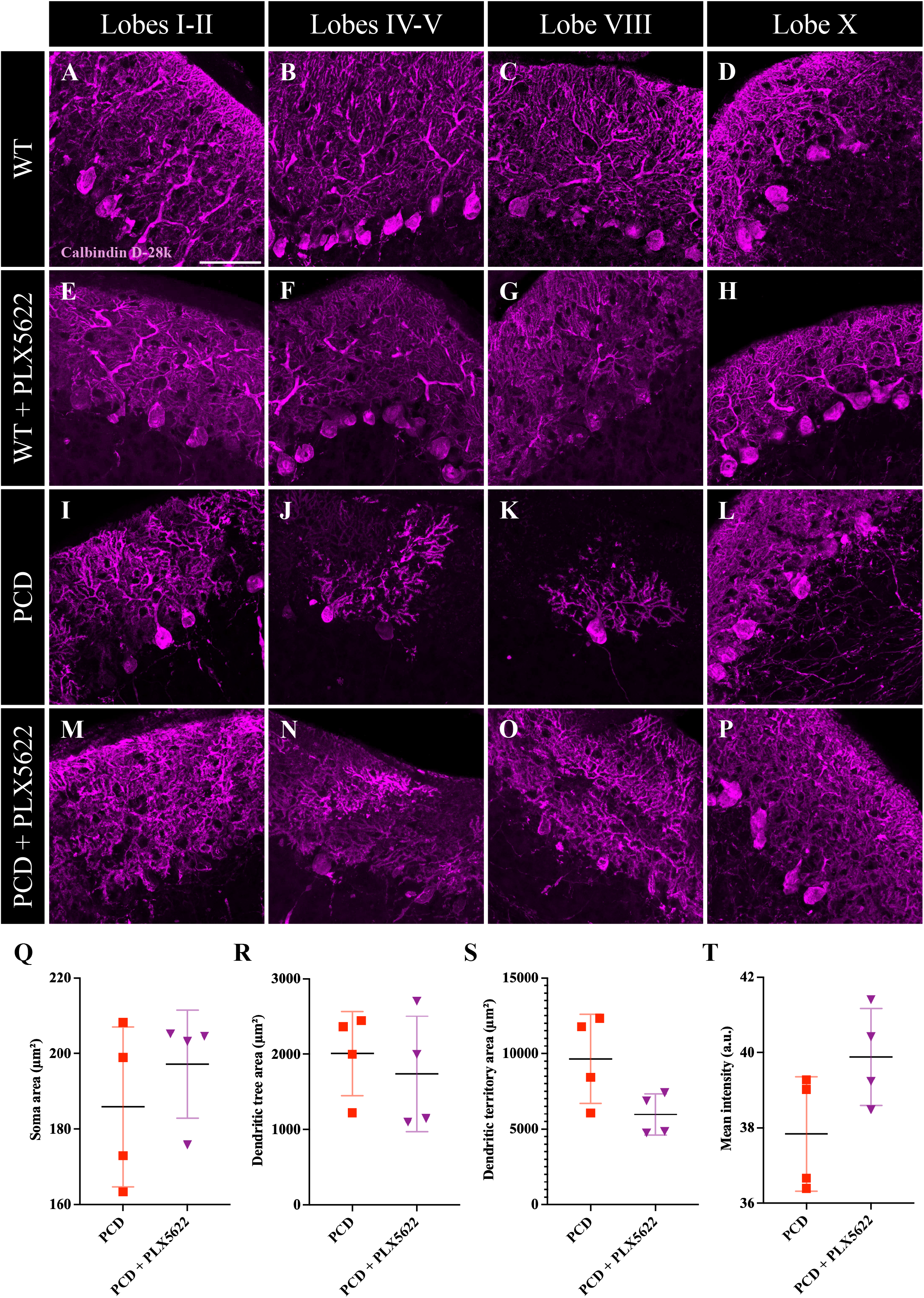
Morphology of surviving Purkinje cells in untreated and PLX5622-treated PCD mice. (A–P) Representative calbindin D-28k immunofluorescence images (magenta) at P25. Rows show WT (A–D), WT + PLX5622 (E–H), PCD (I–L) and PCD + PLX5622 (M–P) mice; columns show cerebellar vermis lobes I–II, IV–V, VIII and X, respectively. (Q–T) Soma area (Q), area occupied by labelled dendritic processes (dendritic tree area; R), area enclosed by the dendritic arbour, including spaces between processes (dendritic territory area; S), and mean calbindin D-28k fluorescence intensity (T). Areas in Q–S are expressed in µmZ. Single-cell quantification was restricted to the two PCD groups because overlapping dendritic labelling prevented reliable segmentation in WT mice, with or without treatment. Each plotted point represents one animal, calculated from the cells analysed across the four lobes. Horizontal black lines and error bars show group means and SD; *n* = 4 independent animals per group. Groups were compared using a two-tailed Mann–Whitney U test; no comparison reached *p* < 0.05. Scale bar, 100 µm. a.u., arbitrary units. WT, wild type; PCD, Purkinje cell degeneration.

Overlapping arbours in the WT groups prevented reliable delineation of individual neurons. Quantitative morphological analysis was therefore restricted to untreated and PLX5622-treated PCD mice, in which sufficiently isolated cells could be selected for segmentation.

Comparison between the PCD groups revealed no significant differences in soma area (*p* = 0.4857; Figure 3Q) or dendritic tree area (*p* = 0.4857; Figure 3R). Dendritic territory area was numerically lower in treated mice, although the difference did not reach statistical significance (*p* = 0.1143; Figure 3S). Mean calbindin D-28k fluorescence intensity was numerically higher after treatment, without a significant between-group difference (*p* = 0.2000; Figure 3T). Comparisons used animal-level means obtained from cells analysed in lobes I–II, IV–V, VIII and X.

Together, greater Purkinje cell preservation at P25 coexisted with neuronal profiles showing qualitative signs of deterioration, without statistically detectable differences between the PCD groups in the measures analysed. Preservation of a larger neuronal population should therefore not be equated with complete structural recovery of all remaining cells. Likewise, the tissue-level increase in calbindin D-28k-positive area described above coexisted with a higher neuronal density, without a significant increase in mean fluorescence intensity per surviving cell.

### PLX5622 reduces cerebellar microglial abundance and attenuates PCD-associated microgliosis

In the representative images, WT mice showed Iba1-positive profiles dispersed throughout the cerebellar tissue. In untreated PCD mice, labelling was more abundant and accumulated particularly in the molecular layer and around the Purkinje cell layer. After PLX5622, labelling appeared less extensive in both genotypes, although Iba1-positive cells remained (Figure 4A–D).

**Figure 4.**
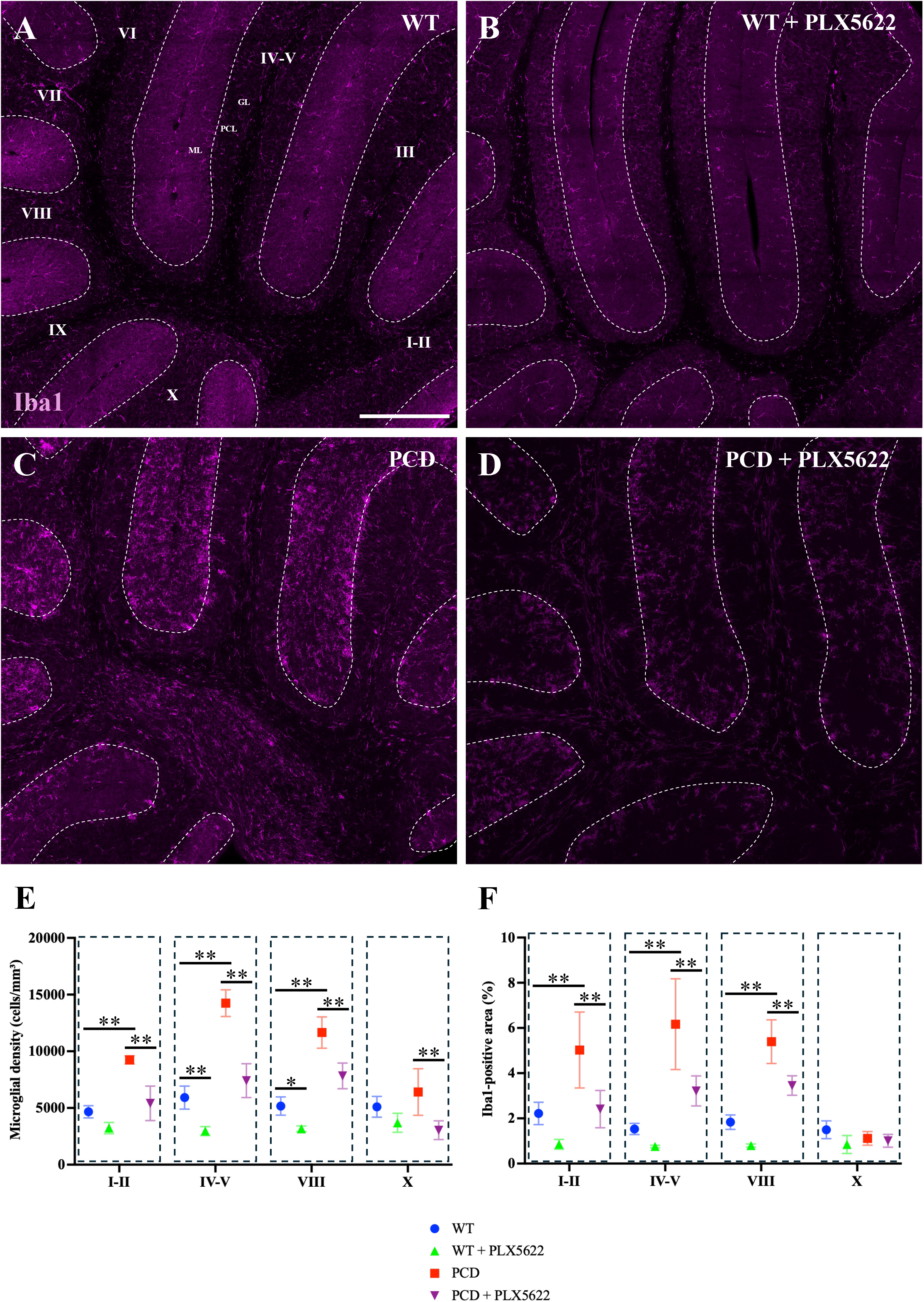
PLX5622 reduces cerebellar microglial abundance and attenuates PCD-associated microgliosis. (A–D) Representative Iba1 immunofluorescence images (magenta) in parasagittal sections of the cerebellar vermis at P25: WT (A), WT + PLX5622 (B), PCD (C) and PCD + PLX5622 (D). Dashed lines indicate the Purkinje cell layer. (E, F) Microglial density (cells/mm3; E) and percentage of the analysed area occupied by Iba1 immunoreactivity (F) in lobes I–II, IV–V, VIII and X. Residual Iba1-positive cells remain visible after treatment. Measurements were summarised within each animal for each lobe. Symbols and error bars represent group means and SD; *n* = 4 independent animals per group. Each lobe was analysed separately using ordinary two-way ANOVA, with genotype and treatment as factors, followed by Bonferroni-adjusted prespecified comparisons (WT versus PCD, WT versus WT + PLX5622, and PCD versus PCD + PLX5622) within each lobe and endpoint. Brackets identify the indicated between-group comparisons; \**p* < 0.05; \*\**p* < 0.01. Scale bar, 200 µm. PCL, Purkinje cell layer; ML, molecular layer; GL, granular layer. WT, wild type; PCD, Purkinje cell degeneration.

Microglial abundance was assessed at P25 by quantifying microglial density and Iba1-positive area in cerebellar vermis lobes I–II, IV–V, VIII and X (Figure 4). Untreated PCD mice had a higher microglial density than WT mice in I–II, IV–V and VIII (all *p* < 0.001), with no significant difference in X (*p* = 0.512). In PCD mice, PLX5622 significantly reduced microglial density in I–II, IV–V and VIII (all *p* < 0.001), as well as in X (*p* = 0.008). In WT mice, treatment reduced this parameter in IV–V (*p* = 0.007) and VIII (*p* = 0.042), whereas the reductions in I–II (*p* = 0.111) and X (*p* = 0.430) did not reach statistical significance. Two-way ANOVA showed significant effects of genotype and treatment in I–II, IV–V and VIII (all *p* < 0.001); in X, only the treatment effect was significant (*p* = 0.003). Genotype × treatment interactions were also observed in I–II (*p* = 0.016) and IV–V (*p* = 0.004), but not in VIII or X (Figure 4E).

Iba1-positive area was significantly increased in untreated PCD mice compared with WT mice in I–II (*p* = 0.0046), IV–V and VIII (both *p* < 0.001), with no significant difference in X (*p* = 0.4405). In PCD mice, PLX5622 significantly reduced Iba1-positive area in I–II (*p* = 0.0076), IV–V (*p* = 0.0060) and VIII (*p* = 0.0010), without a significant change in X (*p* = 1.0000). In WT mice, numerical reductions were observed in all four regions, but none reached statistical significance after adjustment for multiple comparisons: I–II (*p* = 0.2113), IV–V (*p* = 0.9746), VIII (*p* = 0.0635) and X (*p* = 0.0640). ANOVA showed significant effects of genotype in I–II, IV–V and VIII (all *p* < 0.001) and of treatment in I–II (*p* = 0.0015), IV–V (*p* = 0.0044) and VIII (*p* < 0.001). In X, neither genotype (*p* = 0.5422) nor treatment (*p* = 0.0503) had a significant effect, and the genotype × treatment interaction was not significant in any of the lobes analysed (Figure 4F).

Together, PLX5622 attenuated PCD-associated cerebellar microgliosis, with lower microglial density in all four regions and reduced Iba1-positive area in I–II, IV–V and VIII. The persistence of Iba1-positive cells after treatment confirmed that depletion was incomplete. Thus, greater regional Purkinje cell preservation at P25 coexisted with lower microglial abundance and a remaining population of these cells.

### PLX5622 expands the territory encompassed by residual cerebellar microglia

To characterise the microglia remaining after treatment, cell area, soma area, the territory encompassed by processes and integrated Iba1 density were analysed at P25. In representative images, WT microglia showed ramified profiles, whereas untreated PCD microglia had more prominent somata. After PLX5622, cells with extensive processes remained in both genotypes (Figure 5A–P).

**Figure 5.**
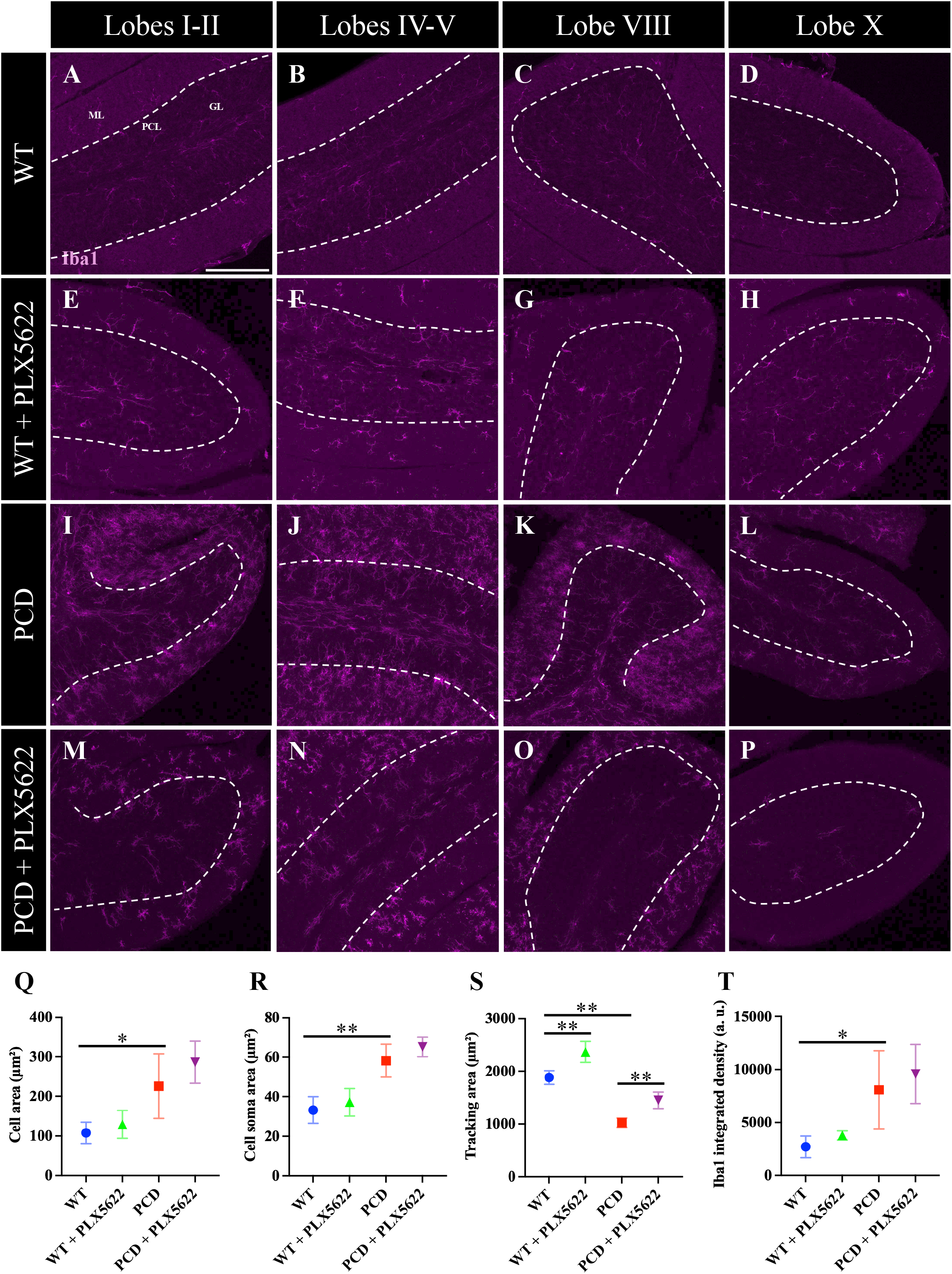
PLX5622 expands the territory encompassed by residual cerebellar microglia. (A–P) Representative Iba1 immunofluorescence images (magenta) at P25. Rows show WT (A–D), WT + PLX5622 (E–H), PCD (I–L) and PCD + PLX5622 (M–P) mice; columns show cerebellar vermis lobes I–II, IV–V, VIII and X, respectively. Dashed lines indicate the Purkinje cell layer. (Q–T) Microglial cell area (Q), soma area (R), area enclosed by the processes, including intervening spaces (tracking area; S), and Iba1 integrated density (T). Areas in Q–S are expressed in µmZ. Tracking area describes the territory encompassed by the processes in the analysed projection. Measurements from the four lobes were combined within each animal before group comparisons. Symbols and error bars represent group means and SD; *n* = 4 independent animals per group. Ordinary two-way ANOVA, with genotype and treatment as factors, was followed by Bonferroni-adjusted prespecified comparisons (WT versus PCD, WT versus WT + PLX5622, and PCD versus PCD + PLX5622). Brackets identify the indicated between-group comparisons; \**p* < 0.05; \*\**p* < 0.01. Scale bar, 100 µm. a.u., arbitrary units; GL, granular layer; ML, molecular layer; PCL, Purkinje cell layer. WT, wild type; PCD, Purkinje cell degeneration.

Quantification showed that untreated PCD microglia had a larger cell area (*p* = 0.026; Figure 5Q) and soma area (*p* < 0.001; Figure 5R) than WT microglia. These differences coexisted with a smaller territory encompassed by their processes (*p* < 0.001; Figure 5S) and higher integrated Iba1 density (*p* = 0.023; Figure 5T).

PLX5622 treatment was accompanied by numerical increases in cell area and soma area in both genotypes, although differences from their respective untreated controls were not significant for cell area (WT, *p* = 1.000; PCD, *p* = 0.410) or soma area (WT, *p* = 1.000; PCD, *p* = 0.522; Figure 5Q, R). By contrast, the territory encompassed by processes increased significantly in both WT (*p* = 0.002) and PCD mice (*p* = 0.005; Figure 5S). Integrated Iba1 density was also numerically higher after treatment in both genotypes, without significant differences from their respective controls (both *p* = 1.000; Figure 5T).

Together, the reduction in microglial density described above was accompanied by an expansion of the territory encompassed by the remaining cells. PLX5622 therefore modified both population abundance and the morphology of residual microglia, with a detectable territorial change in both genotypes.

### PLX5622 regionally reduces CD68 immunoreactivity in PCD mice

In representative images, CD68 labelling was sparse in WT and WT + PLX5622 mice. Untreated PCD mice showed a more extensive signal, with abundant CD68 labelling spatially associated with Iba1-positive profiles, particularly in the molecular layer and around the Purkinje cell layer. In PLX5622-treated PCD mice, labelling was less extensive, although Iba1-positive profiles retaining CD68 signal remained (Figure 6A–L).

**Figure 6.**
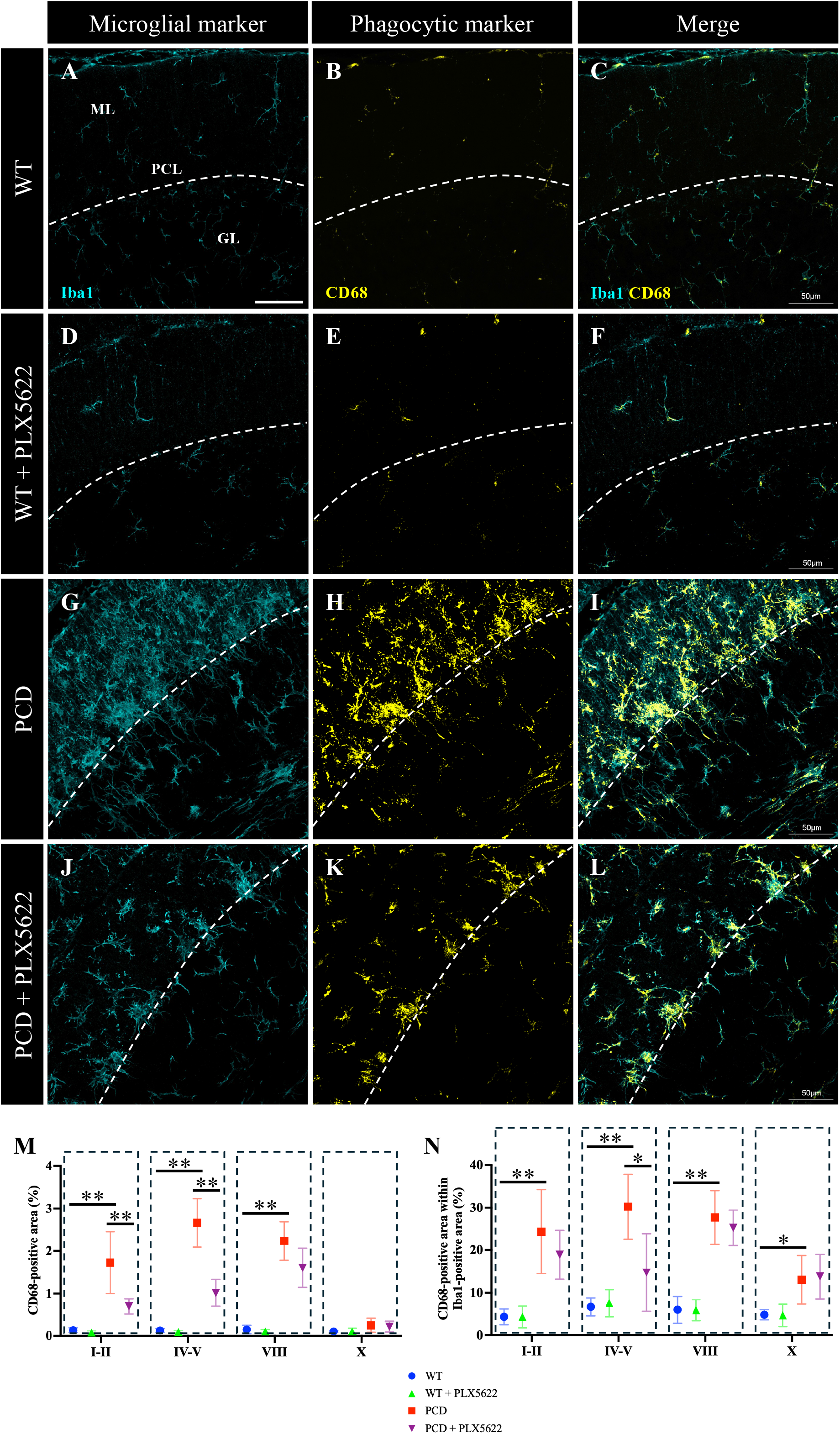
PLX5622 differentially reduces tissue-level and microglia-associated CD68 immunoreactivity in PCD mice. (A–L) Representative confocal images of cerebellar vermis lobes IV–V at P25: WT (A–C), WT + PLX5622 (D–F), PCD (G–I) and PCD + PLX5622 (J–L). Columns show Iba1 (cyan; A, D, G, J), CD68 (yellow; B, E, H, K), and the merged channels (C, F, I, L). Dashed lines indicate the Purkinje cell layer. (M) Percentage of the analysed tissue area within each lobe occupied by CD68-positive staining. (N) Percentage of Iba1-positive area occupied by CD68 immunoreactivity. Both measurements were obtained separately in lobes I–II, IV–V, VIII and X, with one value per animal and lobe. Symbols and error bars represent group means and SD; *n* = 4 independent animals per group. Each lobe was analysed separately using ordinary two-way ANOVA, with genotype and treatment as factors, followed by Bonferroni-adjusted prespecified comparisons (WT versus PCD, WT versus WT + PLX5622, and PCD versus PCD + PLX5622) within each lobe and endpoint. Brackets identify the indicated between-group comparisons; \**p* < 0.05; \*\**p* < 0.01. Scale bars, 50 µm. PCL, Purkinje cell layer; ML, molecular layer; GL, granular layer. WT, wild type; PCD, Purkinje cell degeneration.

CD68 immunoreactivity was quantified as the percentage of cerebellar tissue area positive for this marker and the percentage of Iba1-positive area occupied by CD68 in lobes I–II, IV–V, VIII and X. Untreated PCD mice had a larger total CD68-positive area than WT mice in I–II, IV–V and VIII (all *p* < 0.001), with no significant difference in X (*p* = 0.294; Figure 6M). The percentage of Iba1-positive area occupied by CD68 was also higher in PCD mice in I–II (*p* = 0.001), IV–V and VIII (both *p* < 0.001), as well as in X (*p* = 0.047; Figure 6N).

In WT mice, PLX5622 did not significantly alter either parameter in any of the regions analysed (all *p* = 1.000). In PCD mice, treatment reduced total CD68-positive area in I–II (*p* = 0.007) and IV –V (*p* < 0.001). The decrease observed in VIII did not reach statistical significance (*p* = 0.054), and no significant difference was detected in X (*p* = 1.000; Figure 6M).

When CD68 signal was considered relative to the Iba1-positive compartment, treatment produced a significant reduction only in IV–V (*p* = 0.013). No significant changes were detected in I–II (*p* = 0.659), VIII or X (both *p* = 1.000; Figure 6N).

Together, PLX5622 reduced the tissue extent of CD68 labelling in I–II and IV–V of PCD mice, while a decrease in its relative representation within Iba1-positive area was detected in IV–V. This pattern shows regional attenuation of immunoreactivity associated with the lysosomal compartment, with CD68 signal persisting in the residual microglial population.

### PLX5622 reduces cerebellar CD45, CD16/32 and CD206 immunoreactivity in PCD mice

To complement the regional analyses with an integrated measure of the immunoinflammatory response, the percentages of CD45-, CD16/32- and CD206-positive area measured in lobes I–II, IV –V, VIII and X were averaged within each animal. This yielded a single mean percentage per animal for each marker, summarising immunoreactivity across the cerebellar regions assessed (Figure 7).

**Figure 7.**
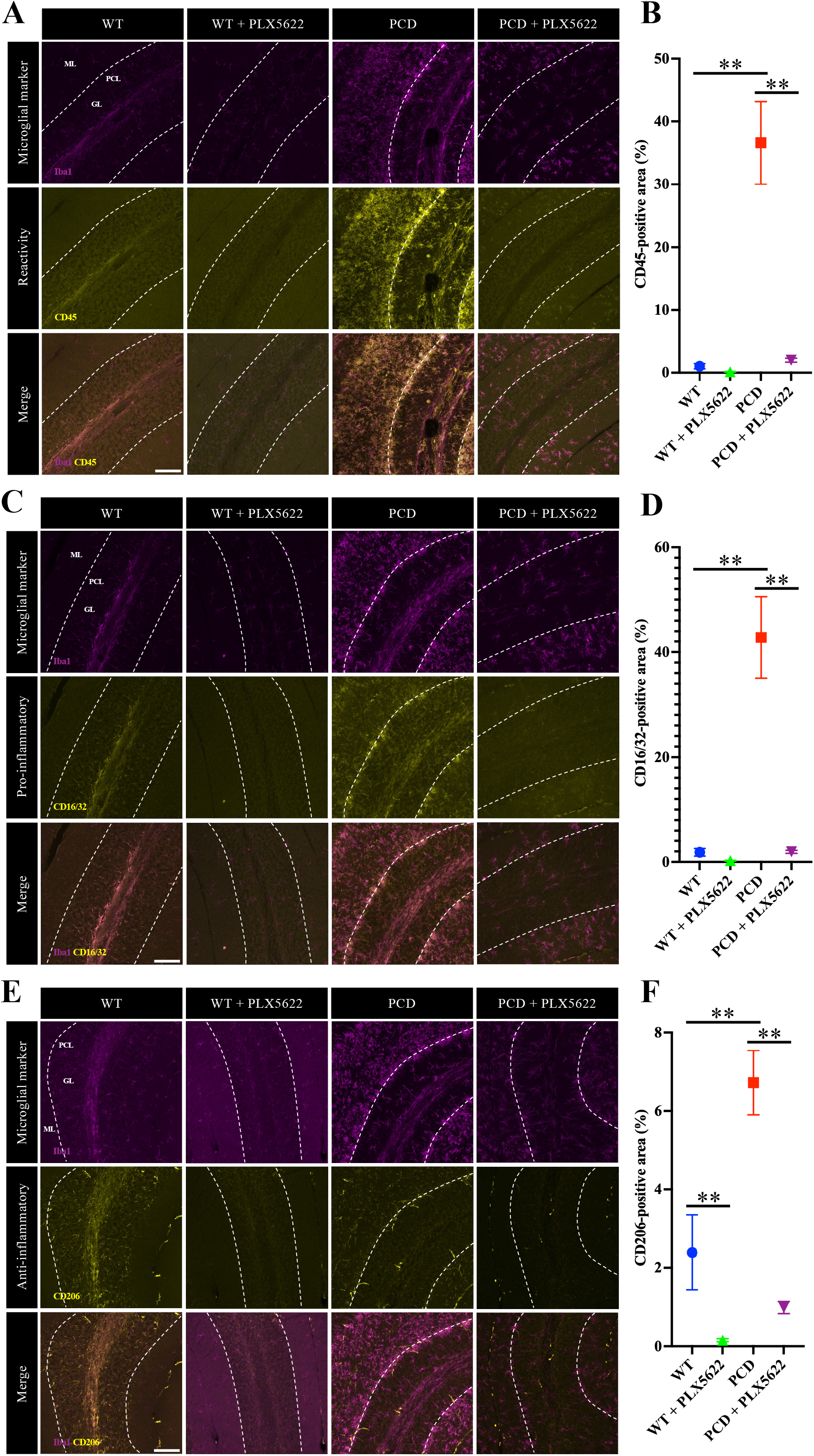
PLX5622 reduces cerebellar CD45, CD16/32 and CD206 immunoreactivity in PCD mice. (A, C, E) Representative immunofluorescence images showing Iba1 (magenta) together with CD45 (A), CD16/32 (C) or CD206 (E) (yellow) at P25. Within each panel, columns show WT, WT + PLX5622, PCD and PCD + PLX5622 mice, from left to right; rows show Iba1, the corresponding marker and the merged channels, from top to bottom. Dashed lines indicate the Purkinje cell layer. (B, D, F) Percentages of analysed area occupied by CD45 (B), CD16/32 (D) or CD206 (F) immunoreactivity. The percentages of positively stained area in lobes I–II, IV–V, VIII and X were averaged within each animal to obtain one mean percentage per marker. Symbols and error bars represent group means and SD; *n* = 4 independent animals per group. Ordinary two-way ANOVA, with genotype and treatment as factors, was followed by Bonferroni-adjusted prespecified comparisons (WT versus PCD, WT versus WT + PLX5622, and PCD versus PCD + PLX5622). Brackets identify the indicated between-group comparisons; \**p* < 0.05; \*\**p* < 0.01. Scale bars, 100 µm. GL, granular layer; ML, molecular layer; PCL, Purkinje cell layer. WT, wild type; PCD, Purkinje cell degeneration.

At P25, untreated PCD mice showed more abundant CD45, CD16/32 and CD206 labelling than WT mice. Qualitative examination indicated reduced labelling of all three markers after PLX5622 in both WT and PCD mice. In merged images, some of this signal was associated with Iba1-positive profiles, and detectable immunoreactivity persisted after treatment in both genotypes (Figure 7A, C, E).

Quantification of positive area showed increased CD45, CD16/32 and CD206 immunoreactivity in untreated PCD mice compared with WT mice (all *p* < 0.001). In PCD mice, PLX5622 significantly reduced positive area for all three markers (all *p* < 0.001; Figure 7B, D, F).

In WT mice, all three markers had lower mean values after treatment, consistent with the qualitative decrease. The reductions in CD45- and CD16/32-positive areas did not reach statistical significance (both *p* = 1.000; Figure 7B, D). By contrast, the reduction in CD206-positive area was significant (*p* < 0.001; Figure 7F).

Descriptively, mean CD45-positive area in treated PCD mice remained above that of untreated WT mice, while CD16/32 showed numerically similar values. Mean CD206-positive area was lower than in untreated WT mice. These observations describe the position of the means and do not represent additional statistical comparisons.

Together, lower microglial abundance was accompanied by reduced CD45, CD16/32 and CD206 labelling in both genotypes, with significant decreases in all three markers in PCD mice. The joint reduction of these signals in the degenerative context shows that treatment attenuates several components of the immune response in cerebellar tissue.

### PLX5622 remodels the expression profile of glial and inflammatory genes in the cerebellum

To complement the histological characterisation, expression of genes associated with microglia, phagocytosis, motility and chemotaxis, and immune and inflammatory responses was analysed by RT–qPCR in whole-cerebellum samples obtained at P25. The heat map summarises the profiles of the four experimental groups relative to untreated WT mice (Figure 8A), while individual analyses are organised into four functional groups (Figure 8B–E).

**Figure 8.**
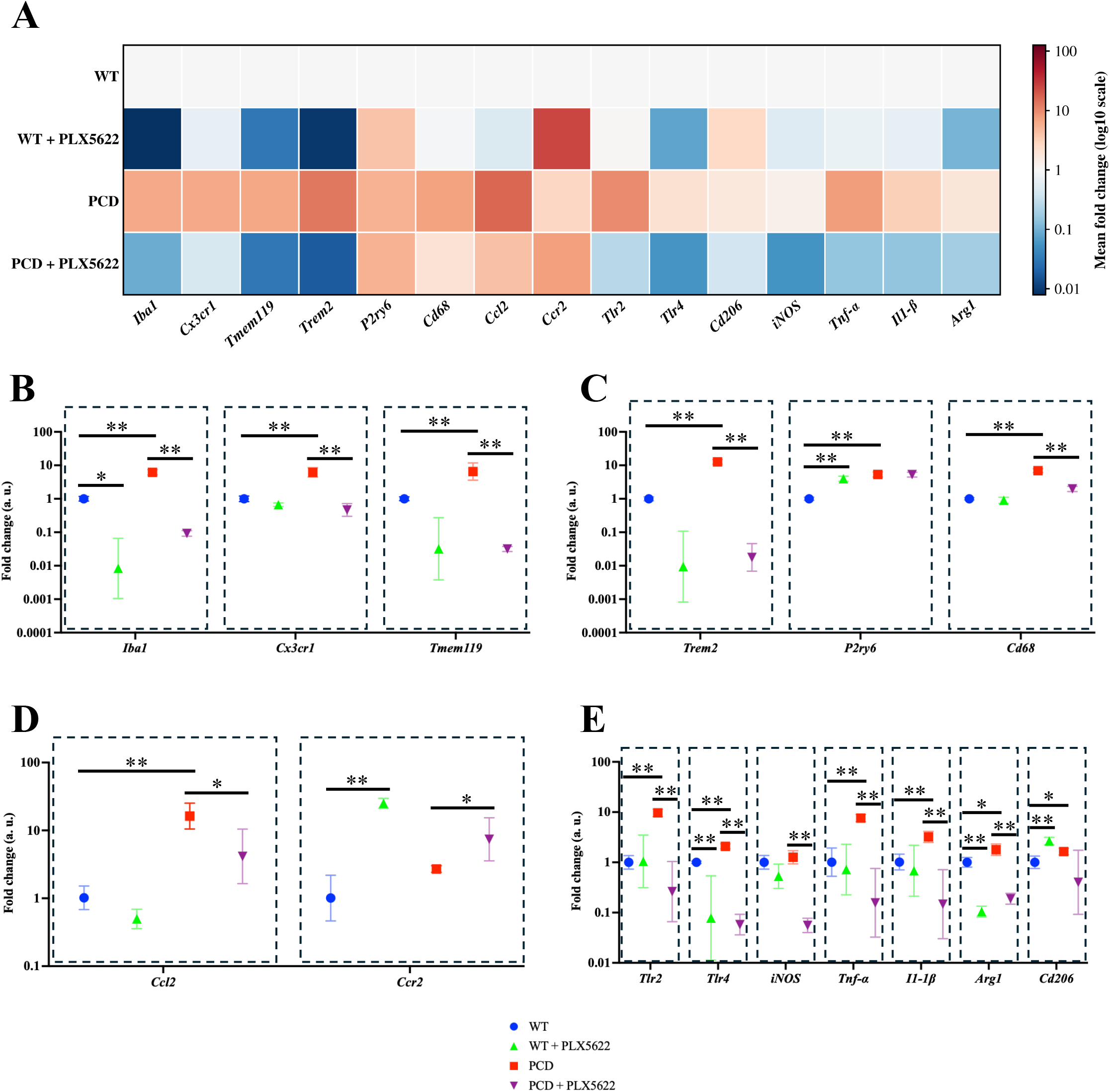
PLX5622 differentially remodels cerebellar gene expression in WT and PCD mice. Gene expression was measured by RT–qPCR in whole-cerebellum samples at P25, normalised to *Gapdh* and expressed as fold change relative to untreated WT mice. (A) Heat map of geometric mean fold changes. The colour scale is logarithmic; red and blue indicate higher and lower expression relative to untreated WT mice, respectively. (B–E) Individual genes organised into four functional groups: microglia-associated markers *Iba1*, *Cx3cr1* and *Tmem119* (B); phagocytosis-associated genes *Trem2*, *P2ry6* and *Cd68* (C); motility- and chemotaxis-associated genes *Ccl2* and *Ccr2* (D); and inflammatory and regulatory genes *Tlr2*, *Tlr4*, *Nos2*, *Tnf*, *Il1b*, *Arg1* and *Cd206* (E). *Nos2*, *Tnf* and *Il1b* are labelled iNOS, Tnf-α and Il-1β, respectively, in the figure. The labels *Iba1* and *Cd206* refer to the mouse genes *Aif1* and *Mrc1*, respectively. Symbols show geometric means, with multiplicative geometric SD error bars on logarithmic y-axes; *n* = 4 independent animals per group. Each gene was analysed using ordinary two-way ANOVA, with genotype and treatment as factors, on untransformed fold-change values. Pairwise comparison and multiplicity procedures are described in Statistical analyses. The displayed contrasts are WT versus PCD, WT versus WT + PLX5622, and PCD versus PCD + PLX5622. Brackets identify the indicated between-group comparisons; \**p* < 0.05; \*\**p* < 0.01. WT, wild type; PCD, Purkinje cell degeneration.

Comparison between untreated mice showed higher expression of the general microglial markers *Iba1*, *Cx3cr1* and *Tmem119* in PCD mice (all *p* < 0.01; Figure 8B). This increase was also observed for the phagocytosis-associated genes *Trem2*, *P2ry6* and *Cd68* (all *p* < 0.01; Figure 8C). Among motility- and chemotaxis-associated genes, PCD mice showed higher *Ccl2* expression (*p* < 0.01), whereas no difference was detected in *Ccr2* (*p* > 0.05; Figure 8D). In the immune and inflammatory response panel, expression of *Tlr2*, *Tlr4*, *Tnf* and *Il1b* increased (all *p* < 0.01), as did *Arg1* (*p* < 0.05), without a significant difference in *Nos2* (iNOS; *p* > 0.05). *Cd206* expression was also higher in PCD mice (*p* < 0.05; Figure 8E).

Examination of PLX5622 effects on general microglial markers showed lower *Iba1* expression in treated WT mice (*p* < 0.05), whereas no significant differences were detected in *Cx3cr1* (*p* > 0.05) or *Tmem119* (*p* > 0.05). In PCD mice, treatment reduced expression of all three markers, *Iba1*, *Cx3cr1* and *Tmem119* (all *p* < 0.01; Figure 8B).

Among phagocytosis-associated genes, PLX5622 increased *P2ry6* expression in WT mice (*p* < 0.01). Mean *Trem2* and *Cd68* values were lower after treatment, although neither difference reached statistical significance (both *p* > 0.05). In PCD mice, treatment significantly reduced *Trem2* and *Cd68* (both *p* < 0.01), while *P2ry6* showed no significant difference (*p* > 0.05; Figure 8C).

In the motility and chemotaxis group, treatment increased *Ccr2* in WT mice (*p* < 0.01), without a significant change in *Ccl2* (*p* > 0.05). In PCD mice, PLX5622 reduced *Ccl2* expression and increased *Ccr2* (both *p* < 0.05; Figure 8D).

Among immune and inflammatory response genes, PLX5622 reduced *Tlr4* and *Arg1* expression and increased *Cd206* in WT mice (all *p* < 0.01), without significant differences in *Tlr2*, *Nos2*, *Tnf* or *Il1b* (all *p* > 0.05). In PCD mice, treatment significantly decreased *Tlr2*, *Tlr4*, *Nos2*, *Tnf*, *Il1b* and *Arg1* (all *p* < 0.01). Mean *Cd206* expression was also lower in treated PCD mice, although this difference did not reach statistical significance (*p* > 0.05; Figure 8E).

To visualise treatment-associated changes within each genotype, the same data were additionally expressed relative to their corresponding untreated control: WT for WT + PLX5622 and PCD for PCD + PLX5622 (Supplementary Figure S1; Additional file 1). This representation complements the WT-referenced profile in Figure 8A and illustrates the predominance of decreases in PCD mice and the combination of selective increases and decreases in WT mice.

Together, PLX5622 was associated in PCD mice with reduced expression of numerous genes across the four functional groups, accompanied by increased *Ccr2* and no significant differences in *P2ry6* or *Cd206*. In WT mice, changes were selective and included both increases and decreases. These results show broad but heterogeneous remodelling of the glial and inflammatory gene-expression profile in cerebellar tissue.

### PLX5622 regionally attenuates the astroglial response in PCD mice

In representative images, GFAP signal was less intense and extensive in WT than in PCD mice. WT mice showed thin, radial processes in the molecular layer and a branched network in the granular layer (Figure 9A). PLX5622-treated WT mice retained this general distribution, although labelling appeared fainter, particularly in the molecular layer (Figure 9B). In PCD mice, processes were more prominent and formed a denser network, particularly in the molecular layer and around the Purkinje cell layer (Figure 9C). After PLX5622 treatment, PCD mice showed less extensive labelling and a less dense network, although GFAP-positive processes persisted in the molecular and granular layers (Figure 9D).

**Figure 9.**
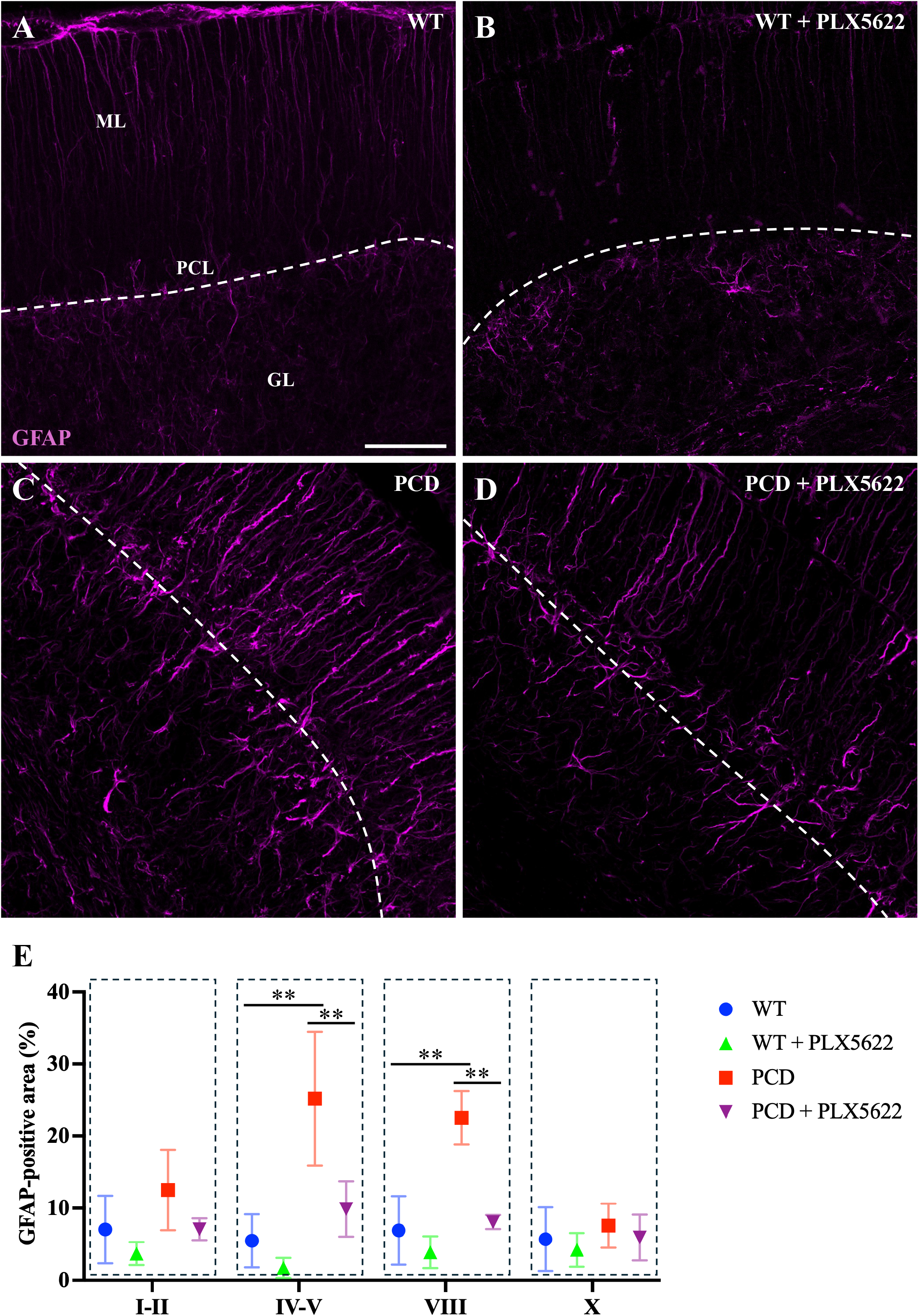
PLX5622 regionally attenuates the cerebellar astroglial response in PCD mice. (A–D) Representative confocal images of GFAP immunoreactivity (magenta) in cerebellar vermis lobe VIII at P25: WT (A), WT + PLX5622 (B), PCD (C) and PCD + PLX5622 (D). Dashed lines indicate the Purkinje cell layer. (E) Percentage of the analysed area occupied by GFAP-positive staining in lobes I–II, IV–V, VIII and X. Measurements were summarised within each animal for each lobe. Symbols and error bars represent group means and SD; *n* = 4 independent animals per group. Each lobe was analysed separately using ordinary two-way ANOVA, with genotype and treatment as factors, followed by Bonferroni-adjusted prespecified comparisons (WT versus PCD, WT versus WT + PLX5622, and PCD versus PCD + PLX5622) within each lobe. Brackets identify the indicated between-group comparisons; \**p* < 0.05; \*\**p* < 0.01. Scale bar, 100 µm. GL, granular layer; ML, molecular layer; PCL, Purkinje cell layer. GFAP, glial fibrillary acidic protein. WT, wild type; PCD, Purkinje cell degeneration.

The astroglial response was assessed by quantifying GFAP-positive area in cerebellar vermis lobes I–II, IV–V, VIII and X at P25 (Figure 9E). Untreated PCD mice had a larger GFAP-positive area than WT mice in lobes IV–V and VIII (both *p* < 0.01), whereas no significant genotype differences were detected in I–II or X (both *p* > 0.05).

After PLX5622 treatment, no statistically significant differences in GFAP-positive area were detected in WT mice in any of the lobes analysed (all *p* > 0.05). In PCD mice, treatment significantly reduced this parameter in IV–V and VIII (both *p* < 0.01), without significant differences in I–II or X (both *p* > 0.05). ANOVA showed a significant genotype × treatment interaction in lobe VIII (*p* < 0.01), whereas this interaction did not reach significance in I–II, IV–V or X (all *p* > 0.05).

Together, significant reductions in GFAP immunoreactivity in treated PCD mice were detected in the two territories showing a significant increase in this marker relative to WT mice. These results support regional attenuation of the astroglial response and extend the treatment-associated changes in the cerebellar environment to the astroglial component.

## Discussion

Using the early PLX5622 regimen established in our previous study [41], we examined when the behavioural response emerged and which regional glial changes accompanied Purkinje cell preservation. In PCD mice, improved motor coordination was detected at P25, alongside improvements in selected spontaneous behaviours and novel-object discrimination. At that age, microglial density was lower in all four PCD regions, while greater Purkinje cell preservation and lower GFAP immunoreactivity coincided in lobes IV–V and VIII. The territory encompassed by the processes of residual microglia expanded in both genotypes, and the fraction of Iba1-positive area labelled for CD68 decreased in lobes IV–V in PCD mice. CD45, CD16/32 and CD206 immunoreactivity decreased in PCD mice. The transcriptional response was selective, with several inflammatory transcripts decreasing in PCD mice and selected transcripts increasing in WT mice.

Significant improvements in rotarod performance were detected at P25, but not at P17 or P21. At P25, PCD mice undergo intense Purkinje cell loss [41, 72, 73], so preserving a larger neuronal population could have a more evident motor impact. At earlier ages, less pronounced impairment could leave less scope to detect this benefit. Purkinje cells constitute the output of the cerebellar cortex [19], and their preservation could help maintain coordination, as suggested by the functional benefits associated with neuroprotection in other PCD studies [61, 74]. Microglial signals can also modify neuronal excitability during cerebellar inflammation [27], providing another potential contribution to the functional changes. Treatment began at P15, when our previous work had identified altered expression of microglial and inflammatory genes before overt Purkinje cell death and without a detectable increase in microglial density [41]. The early molecular response therefore precedes the marked histological microgliosis accompanying neuronal loss. Intervention during this interval could limit responses that subsequently amplify damage, helping explain why a benefit becomes evident when degeneration is more advanced. This sequence supports investigating early molecular changes as possible indicators of an intervention window, while neurons susceptible to preservation remain.

The spontaneous-behaviour and recognition results extend this functional assessment beyond rotarod performance. At P25, treated PCD mice spent more time grooming and walking and less time resting, whereas rearing did not improve significantly. The response was therefore selective, rather than a uniform recovery of spontaneous activity. Novel-object discrimination also improved without a significant change in total exploration time, making a simple increase in the time available for exploration a less sufficient explanation. The exploration percentage and discrimination index describe the same recognition outcome and should be interpreted together. Cognitive benefits after partial CSF1R inhibition have also been reported in 3xTg-AD mice, although that study found improved place recognition without improved object recognition [75].

This difference supports considering the particular task and pathological context when interpreting the benefit. In our mice, the combination of anatomical preservation and improvements across several behavioural measures is consistent with better functional maintenance during degeneration; systemic treatment and the involvement of distributed circuits mean that the recognition effect cannot be assigned exclusively to the cerebellum.

The early behavioural changes in WT mice provide a complementary perspective on this context dependence. PLX5622 reduced grooming across the ages assessed, while changes in rearing, walking, resting and novel-object discrimination were more evident at earlier assessments. Early exposure to PLX5622 can therefore affect behaviour during development in WT mice. In adult WT mice, PLX5622 did not impair open-field or Barnes-maze performance [75], whereas CSF-1-dependent cerebellar microglia have been implicated in motor function and social interaction [26]. Age, exposure and the behavioural domain examined may contribute to the different outcomes. Physiological microglia–neuron interactions also provide a biological basis for functional effects outside injury [76], while exploratory behaviour remains sensitive to stress and testing conditions [77]. These precedents favour interpreting the WT changes as part of the response to intervention in a developing nervous system. The distinct trajectory in PCD is compatible with an additional benefit from limiting an injury-associated response.

Previous studies from our group have shown that PCD degeneration can be modified by interventions with behavioural benefits. Oleoylethanolamide (OEA) improved motor, cognitive and social functions [60] and, in a complementary study, reduced microgliosis and modified the inflammatory response [42]. Recombinant human vascular endothelial growth factor B (rhVEGF-B) also produced neuroprotection and behavioural benefits [61]. These studies relate neuronal preservation to function, although the contributions of direct neuronal protection and changes in the inflammatory environment are difficult to separate. Our initial PLX5622 study reported Purkinje cell preservation and motor improvement at P25 [41]. The larger rotarod cohort and additional behavioural measures in the present study extend that functional evidence. The reduction in microglial density was consistent with the intended action of PLX5622, and systemic CSF1R inhibition can also affect other immune compartments [52, 53].

Motor improvement was accompanied by greater Purkinje cell preservation. At P25, PLX5622 was associated with higher neuronal density and a larger calbindin-positive area in IV–V and VIII, whereas no significant preservation was detected in I–II or X. Cerebellar lobes differ in their molecular organisation and vulnerability [36, 37], and the anterior regions are more affected in PCD, whereas X retains its neurons for longer [37, 72, 73]. Purkinje cell loss is rapid and asynchronous, and surviving cells are concentrated in X at P30 [72]. In another allele of the model, loss affected I–VIII between P21 and P30, whereas some neurons in IX–X persisted for additional weeks [73]. Assessment of all territories at P25 may therefore capture different degenerative stages within the same cerebellum. This temporal difference could help explain the distribution of protection. In I–II, more advanced damage could reduce the scope for preservation, whereas IV–V and VIII would retain a population more susceptible to benefiting from the intervention. The situation in X is different: in studies from our group, neuronal density was maintained until P30 and declined at P35 [37]. Its resistance, related to specific glutamylation patterns and to heat shock protein 25 (HSP25) and its phosphorylation at Ser15 [36, 37], could leave less scope to detect an additional benefit at P25. These possibilities relate the treatment response to the local tissue state, without establishing a strict sequence of degeneration across all lobes.

Greater Purkinje cell preservation coexisted with heterogeneous neuronal morphology. Some surviving cells had restricted dendritic arbours and apparently small somata, but none of the quantified morphological parameters differed significantly between the PCD groups. These observations describe the condition of individual surviving neurons; they do not establish treatment-induced soma shrinkage, and the mean soma area was numerically higher in treated mice. Healthy Purkinje cells have an extensive, branched and predominantly parasagittal arbour that integrates climbing and parallel fibre inputs [78, 79], with dendritic reorganisation still ongoing around P25 [78]. This anatomical reference helps interpret the restricted profiles observed in PCD.

Individual WT morphometric values could not be obtained because the densely labelled arbours overlapped, a segmentation difficulty also described previously [79]. The comparison therefore concerns the surviving PCD population rather than a measured return to WT dimensions. CCP1-related dendritic abnormalities described in another allele could persist despite reduced injury from the surrounding glial environment [73]. Likewise, greater neuronal preservation after rhVEGF-B in our model coexisted with apoptotic activation [61]. PLX5622 may thus favour the persistence of vulnerable neurons without producing complete structural recovery at the age examined.

PLX5622 reduced microglial density in all four PCD territories and produced significant reductions in IV–V and VIII in WT mice. The effect in WT mice is consistent with the dependence of microglial survival on CSF1R under physiological conditions and with the relationship between PLX5622 concentration and the degree of depletion [75]. In PCD, treatment additionally acts on a population expanded during degeneration [41]. CSF1R also regulates proliferation: in APP/PS1 mice, its inhibition with GW2580 limited pathology-associated microglial expansion [80]. Lower density in PCD could thus result from both cell elimination and reduced population expansion during injury. CCP1 deficiency could also influence the response to PLX5622. CCP1 participates in tubulin deglutamylation, and its absence is associated with brain hyperglutamylation [81]. Our previous work showed functional alterations in microglial cells deficient in this enzyme [41]. The importance of microtubules in the microglial response has been demonstrated following lipopolysaccharide (LPS) stimulation, when their reorganisation contributes to morphological changes and cytokine transport and release [82]. Moreover, in a macrophage cell line, experimental microtubule disorganisation altered the formation and distribution of CSF1R signalling complexes [83]. A microtubule alteration could therefore modify the response to inhibition of this receptor. This hypothesis links the CCP1 defect to the treatment target, but does not demonstrate increased sensitivity of PCD microglia to PLX5622. Reducing this population could limit its participation in mechanisms that aggravate primary neuronal vulnerability. In prion disease, inhibition of proliferation with GW2580 reduced neuronal injury and delayed progression [84]; in PCD, minocycline delayed Purkinje cell death and produced a mild motor improvement [74]. Together with our previous PLX5622 findings [41], these results support the possibility that injury-associated responses can modify neuronal loss. Furthermore, complete elimination does not appear necessary to obtain a benefit: improved spatial learning and place recognition were reported with partial depletion in 3xTg-AD mice [75]. Similarly, early treatment with PLX3397 in a mouse model of spinocerebellar ataxia type 1 (SCA1) reduced cerebellar microglial density and *Aif1* and TNFα expression and improved motor performance despite incomplete depletion [85]. In our study, more neurons were also preserved despite the persistence of a residual microglial population, making its characteristics relevant to understanding the treatment effect.

Microglial density and Iba1-positive area provided complementary measures of this reduction. In WT mice, density decreased significantly in IV–V and VIII, while the numerical decreases in Iba1-positive area did not reach significance. Density reflects the number of cells within the sampled volume, whereas positive area also depends on labelled somata and processes, antigen distribution and the segmentation threshold. A lower cell count need not therefore produce a proportional decrease in labelled area. At P25, our previous study found larger somata and smaller territories encompassed by processes in cerebellar PCD microglia than in WT microglia [41]. Here, the territory encompassed by the processes expanded significantly after PLX5622 in both genotypes. The numerical increases in cell area, soma area and integrated Iba1 density did not reach statistical significance. Qu et al. also described hypertrophic residual microglia in both WT and SCA1 mice after PLX3397 [85], providing a qualitative parallel to the remodelling observed here. After PLX5622 in rats with spinal cord injury, striatal arborisation increased without a significant soma enlargement, and perilesional soma area also increased numerically without significance [86]. A similar non-significant soma increase was reported after PLX3397 in adult mice, although process ramification decreased in that study [87]. In 3xTg-AD mice, partial PLX5622 depletion instead produced a significant increase in residual cell size together with lower Iba1 staining intensity [75]. These findings support considering morphological remodelling and changes in labelling when interpreting density and area together, while showing that the precise response varies across paradigms. The shared increase in territory is compatible with a response to lower cell density in both genotypes. The territory enclosed by the processes is nevertheless distinct from the area of Iba1-positive pixels, and neither the numerical enlargement nor the lack of a significant tissue-area change demonstrates functional compensation. With four animals per group, the pooled residual variability estimated from all four groups and the multiple-comparison adjustment also influence which differences are detected. Consistent with this distinction, ramified residual cells in the visual cortex can coexist with reduced overall tissue surveillance during depletion [88]. Thus, a larger individual territory can accompany a substantial reduction in the collective microglial response.

The CD68 results further distinguish the extent of the tissue response from the characteristics of the residual population. In PCD, CD68-positive tissue area decreased significantly in I–II and IV–V, whereas its relative representation within Iba1-positive area decreased significantly only in IV–V. The latter measure describes the proportion of the labelled microglial compartment occupied by CD68; it is influenced by both labels and is not a measure of phagocytic uptake per cell. Residual microglia can show different lysosomal responses after CSF1R inhibition: the proportion of CD68-positive microglia decreased after short-term PLX5622 in 5xFAD mice [51], but increased in knock-in *App* models [89]. In healthy adult mice, PLX3397 also increased CD68 in isolated residual microglia despite depletion [87]. These measurements differ from our area analysis, but demonstrate why population reduction need not suppress lysosomal markers uniformly in the cells that remain. Our previous functional assays similarly showed that the PCD phenotype depends on cellular origin and environment. Bone marrow-derived microglia-like cells internalised fewer microspheres, while cortical microglia internalised more even after accounting for cell area; cerebellar microglia took up more neuronal debris, but the difference was not significant after area normalisation [41]. The current tissue measurements complement those assays by showing how the extent of a lysosomal response changes during treatment, rather than directly quantifying the same process. The early increases in *Trem2*, *Dap12* and *P2ry6* at P15, followed by increased *Cd68* during degeneration [41], also place the present decreases in *Trem2* and *Cd68* within a previously characterised temporal response. The absence of a significant *P2ry6* decrease shows that this attenuation is selective. Studies separating phagocytic uptake from intracellular lipid handling further illustrate why lysosomal labelling cannot by itself define the efficiency or consequences of clearance [90].

The regional pattern may reflect how the remaining cells encounter injury signals. Apoptotic material can induce clearance-related transcriptional programmes in isolated forebrain microglia [91], and neuronal apoptosis in the developing retina promotes lysosome-enriched microglial states with altered dependence on CSF1R signalling [92]. In IV–V, greater neuronal preservation together with lower total and relative CD68 labelling is compatible with a lower burden of material stimulating this response. In I–II, a tissue-area decrease without a significant change relative to Iba1 is compatible with a stronger contribution from reduced cellular abundance. In VIII, preservation of neuronal somata can coexist with ongoing dendritic or other subcellular damage: microglial interactions and dendritic engulfment before soma loss have been documented in another model of Purkinje cell degeneration [93]. In lobe X, where Purkinje cell loss occurs later [37, 73], CD68 occupied a larger fraction of Iba1-positive area in untreated PCD mice at P25, when genotype differences in microglial density and total CD68-positive area were undetectable. Our previous study detected a rise in microglial density in X at P40 [41]. Relative CD68 labelling can therefore reveal a local difference at a stage when a difference in cell density is still undetectable. These possibilities explain why the same treatment need not yield identical CD68 patterns across regions, although differences in significance do not establish a difference between regional treatment effects. The persistence of CD68-positive profiles near the Purkinje cell layer and the heterogeneous condition of surviving Purkinje cells are compatible with ongoing local responses to injury during treatment. A smaller microglial population could contribute less to neuronal injury while some residual cells retain lysosomal CD68 labelling.

The combined CD45, CD16/32 and CD206 analysis addressed the integrated immune response in the same cerebellar regions examined separately for neuronal and microglial measures. Averaging the regional area percentages within each animal provides a complementary view of mean immunoreactivity across the sampled regions. This biological perspective is relevant because signals released by microglia can influence neighbouring astrocytes and sustain intercellular responses [43]. Such communication does not imply uniform inflammation: a common systemic LPS stimulus can elicit different microglial and astrocytic responses across brain regions [94]. Our integrated measurement therefore describes the sampled regions collectively, while the regional analyses retain information about local vulnerability. At P25, we previously found higher integrated CD45 fluorescence and larger proportions of Iba1-positive cells labelled for CD16/32 or CD206 in the PCD cerebellum [41]. Here, PLX5622 reduced the tissue area positive for each of the three markers in PCD mice. This decrease describes the overall extent of labelling across the sampled regions. In WT mice, CD45 and CD16/32 decreased numerically without significance, whereas CD206 decreased significantly. Fewer cells contributing to the signal probably account for part of this tissue-level attenuation. PLX3397 reduced cerebral CD45-positive populations [95], and PLX5622 after status epilepticus reduced microglial abundance and inflammatory mediators without a detectable change in the relative expression of CD16/32 and CD206 [96]. The cellular source is particularly relevant to CD206: CSF1R inhibition can also reduce CD206-positive perivascular macrophages [97, 98]. Its decrease in WT mice may therefore include effects on populations present without degeneration. Changes in the remaining cells are another possible contribution. In experimental autoimmune encephalomyelitis, PLX5622 reduced the iNOS-positive fraction of Iba1-positive cells while the Arg1-positive fraction was unchanged in most comparisons [99]. Together with the heterogeneity identified by single-cell analyses [100], these findings support interpreting the joint decrease as attenuation of several components of the injury-associated immune response. A reduction in markers associated with different functions need not represent a uniform change of state or a specific loss of reparative capacity.

The molecular results extend this interpretation across four functional groups. Among general microglial markers, *Iba1*, *Cx3cr1* and *Tmem119* decreased significantly in PCD, consistent with a smaller contribution from the population that had expanded during degeneration [41]. In our previous time course, cerebellar microglial density remained elevated at P40, when fewer of the assayed transcripts were still elevated [41]. In WT mice, *Iba1* decreased, while the numerical decreases in *Cx3cr1* and *Tmem119* were not significant. The transcriptional response therefore did not simply reproduce an identical reduction across markers and genotypes. After spinal cord injury, PLX5622 reduced microglial transcripts in both sham-operated and injured mice, alongside suppression of an injury-related inflammatory programme [49]. Moreover, isolated microglia surviving PLX3397 in healthy adults downregulated homeostatic genes while upregulating selected immune and lysosomal genes [87]. These precedents identify two processes that can coexist during depletion: changes in the number and composition of cells contributing RNA, and regulation within the surviving cells. In our whole-cerebellum samples, both may contribute to the net expression measured. Their relative importance can differ between WT tissue undergoing postnatal development and the PCD cerebellum, where a strong degenerative response is already present.

Among phagocytosis-associated genes, *Trem2* and *Cd68* decreased significantly in PCD, while *P2ry6* showed no significant change. In WT mice, *P2ry6* increased, whereas the numerical reductions in *Trem2* and *Cd68* were not significant. The *P2ry6* result is therefore a selective WT increase rather than a demonstrated opposite-direction effect in PCD. PLX5622 reduced *Trem2* in both WT and APP-PS1 mice in a longer adult treatment paradigm [101], showing that the absence of a significant WT reduction in our study is not an invariant response to CSF1R inhibition. Conversely, the increased lysosomal and immune signatures of isolated residual cells after PLX3397 show that population loss can coexist with enhanced expression of selected functional components [87]. Responses to material requiring clearance could contribute to such regulation [91], providing a plausible context for the WT *P2ry6* increase, although neither its cellular source nor increased phagocytic uptake is established by this transcript. In PCD, reduced *Trem2* and *Cd68* accompany the lower tissue CD68 burden, while the absence of a significant *P2ry6* decrease and residual CD68-positive profiles indicate that the response is remodelled rather than uniformly abolished.

The motility and chemotaxis group showed another selective response: *Ccl2* decreased in PCD, while *Ccr2* increased in both genotypes. Our previous study had identified increased *Ccl2* before overt Purkinje cell loss and greater motility in microglia isolated from the cerebellum at P25 [41], so the present reduction is consistent with attenuation of a previously established chemotactic component. The concurrent *Ccr2* increase nevertheless argues against treating ligand and receptor expression as a single uniformly suppressed pathway. CCR2 is strongly associated with recruited monocytes rather than being a selective readout of resident microglial motility [102]. Increased accumulation of CCR2-positive macrophages after PLX5622 has been demonstrated in cerebral ischaemia [103], providing a precedent for a reduced microglial population coexisting with greater representation of another immune population. This does not demonstrate recruitment in our mice. It does, however, support interpreting bulk *Ccr2* expression in relation to cellular composition and the different tissue environments, rather than assuming that the remaining microglia became more motile. The shared direction of the *Ccr2* change also shows that the WT and PCD responses are not opposites for every gene.

Among immune-response genes, the decreases in *Tlr2*, *Tlr4*, *Tnf*, *Il1b* and *Nos2* in PCD indicate attenuation of components associated with danger recognition and inflammatory signalling. Several of these genes were already elevated during degeneration in our previous study [41]. Lower proinflammatory cytokine expression has also followed PLX5622 treatment after status epilepticus [96], while PLX3397 reduced LPS-induced cerebral TNFα and IL-1β without changing their basal concentrations [95]. These findings support considering the pre-existing inflammatory burden when interpreting the more extensive decreases in PCD. Within the same panel, *Arg1* decreased and *Cd206* showed a numerical but non-significant reduction in treated PCD mice. This is compatible with attenuation of a mixed injury-associated response that includes regulatory components. In WT mice, increased *Cd206* coexisted with decreased *Arg1* and *Tlr4*, again arguing against a uniform shift between opposing functional categories. *Cd206* expression also changed in different directions under different irradiation conditions after transient PLX5622 depletion and repopulation [104], although that later phase differs from our assessment at the end of the treatment course. The relevance of context is further illustrated by adult APP-PS1 mice, where depletion increased cortical *Ccl2* and *Il1b* rather than suppressing them [101]. The literature therefore supports selective and disease-dependent responses, rather than an obligatory anti-inflammatory effect of depletion. In our WT mice, higher *Cd206* RNA and a smaller CD206-positive tissue area can coexist because relative transcript abundance and the spatial distribution of protein measure different aspects of the response. Taken together, the WT increases in *P2ry6*, *Ccr2* and *Cd206* are compatible with selective regulation and altered cellular contributions during depletion, while in PCD these processes occur against an expanded degenerative response whose tissue and molecular burden is reduced.

GFAP immunoreactivity decreased in IV–V and VIII in treated PCD mice, indicating attenuation of this component of the astroglial reaction. Our group had described increased GFAP, process hypertrophy and disorganisation of Bergmann fibres during Purkinje cell loss [39]. PLX5622 modifies a response that accompanies degeneration in this model. Similar effects have been observed with PLX5622 following status epilepticus [96], during α-synuclein overexpression [105] and in experimental sepsis, even with pretreatment that produced partial depletion [106]. Likewise, PLX3397 reduced the signal from a GFAP reporter during follow-up of Tg2541 tauopathy [107]. One possible explanation is that a smaller microglial population provides fewer inflammatory stimuli to astrocytes. Conditioned-medium experiments show that signals released by activated microglia can induce reactive responses in these cells [43]. Furthermore, in Tg2541, microglia persisting after CSF1R inhibition exhibited different morphological and molecular profiles according to experimental conditions [107]. In our PCD mice, reduced microglial density and lower abundance of inflammatory transcripts are compatible with weaker microglial stimulation of astrocytes. Greater Purkinje cell preservation could also contribute by reducing the injury signals that maintain the astroglial reaction. Both possibilities may act together: as the microglial contribution to damage decreases, more neurons would be preserved and the stimuli sustaining gliosis would be limited. If treatment delays components of degeneration, the lower GFAP signal at P25 could also reflect a change in the course of this response [39]. Reduced labelling may also result from changes in GFAP expression or in the extent of the processes containing it. After helium irradiation, PLX5622 reduced both immunoreactivity and GFAP-positive volume per cell [108]. Our result may therefore reflect less expansion of astroglial labelling without implying astrocyte loss.

The absence of a significant GFAP decrease in I–II and X can be considered in the context of their different degenerative stages. Bergmann glia have a zonal molecular organisation related to Purkinje cell compartmentalisation [109], and glial responses to a shared inflammatory stimulus can differ regionally [94]. In I–II, ongoing injury and a sustained astroglial response could maintain GFAP despite fewer microglia; reactive astroglial programmes can persist after removal of their inducing stimuli in culture [43]. In X, the more prolonged preservation of Purkinje cells [37] provides a different injury burden and less scope for detecting the same response at P25. The lack of significant baseline GFAP elevations in I–II and X also distinguishes them from IV–V and VIII, where both the PCD-associated increase and the treatment-associated decrease were detected. Microglial reduction does not invariably reduce astrogliosis: partial PLX5622 depletion in 3xTg-AD mice left the plaque-associated astroglial reaction unchanged [75]. Similarly, the SCA1 study detected no significant improvement in GFAP intensity, calbindin intensity or molecular layer width despite better motor performance [85]. Qu et al. administered PLX3397 in chow from weaning for approximately two months [85], whereas we administered PLX5622 by intraperitoneal injection from P15 to P24. Differences in compound, administration route and treatment duration should therefore be considered when comparing the glial responses observed in the two studies. Thus, the coincidence of lower GFAP and greater Purkinje cell preservation in IV–V and VIII is compatible with a joint contribution from reduced microglial stimulation and lower neuronal injury, shaped by local tissue conditions.

The results support the possibility that the microglial response amplifies damage initiated by CCP1 deficiency. Greater Purkinje cell preservation, despite coexisting structural deterioration, suggests that it is possible to favour the persistence of neurons that remain exposed to the genetic defect. The protection obtained by modifying the glial environment identifies a component of degeneration that can be targeted to preserve part of the neuronal population and its function. The rapid Purkinje cell degeneration and intense microglial response developing in PCD [39, 41] make it particularly interesting to explore interventions restricted to phases in which these cells contribute to amplifying damage. In experimental traumatic brain injury, a delayed seven-day course of PLX5622 during the chronic phase, followed by withdrawal and repopulation, reduced neurodegeneration and improved subsequent functional outcomes [48]. In another study, turnover and repopulation through pharmacological or genetic approaches promoted repair and cognitive recovery, whereas microglial elimination alone had limited effects [110]. Thus, the value of a transient intervention may also lie in subsequently allowing microglia with reparative functions to participate. However, its application must consider the functions performed by microglia at each phase of injury. In a hippocampal lesion model, microglial depletion with PLX3397 during lesion induction aggravated neuronal loss, whereas post-lesion depletion improved functional recovery [111]. Likewise, in independent prion disease studies, inhibition of proliferation with GW2580 delayed progression [84], whereas extensive depletion with PLX5622 accelerated it [112]. These differences show the importance of preserving protective functions when designing an intervention targeting the microglial population. Our findings provide a preclinical basis for investigating this possibility at defined disease stages. The aim would be to identify when microglia contribute to damage and to adjust the onset, intensity and duration of treatment accordingly. Transient depletion followed by repopulation is one of the strategies that could be explored [48, 110]. Its potential will depend on limiting harmful contributions while preserving or supporting the functions required for tissue maintenance and repair [84, 112].

## Conclusions

In this study, we show that early PLX5622 treatment is accompanied by greater regional Purkinje cell preservation and functional improvement at P25 in PCD mice. These benefits coexisted with a reduction in the microglial population, morphological changes in the remaining cells and selective attenuation of cerebellar glial and inflammatory responses. Our findings therefore support a contribution of the neuroimmune response to the amplification of neuronal injury in this model and suggest that its modulation can help preserve neurons that remain exposed to the primary genetic defect. The protection achieved with incomplete depletion also shows that the benefit obtained with this regimen does not require complete microglial elimination. The residual population and the associated astroglial changes are thus particularly relevant to understanding the conditions accompanying neuronal preservation. However, differences between cerebellar regions and the distinct behavioural responses of WT and PCD mice indicate that the consequences of this intervention depend on the pathological and temporal context. These findings support investigating interventions that modulate microglial responses during defined phases of degeneration, adjusting their intensity and duration to limit processes that aggravate injury while retaining the functions required for tissue homeostasis and repair.

This work reinforces the glial and inflammatory environment as a modifiable component of PCD neurodegeneration. Targeting this environment offers scope for neuronal and functional protection even when the genetic cause of disease persists, and supports modulation of the microglial response as a complementary approach to limiting its consequences.

## List of abbreviations

ANOVA: analysis of variance;
CCP1: cytosolic carboxypeptidase 1;
CONDCA: childhood-onset neurodegeneration with cerebellar atrophy;
CSF1R: colony-stimulating factor 1 receptor;
DAPI: 4′,6-diamidino-2-phenylindole;
DI: discrimination index;
GFAP: glial fibrillary acidic protein;
GL: granular layer;
HSP25: heat shock protein 25;
Iba1: ionised calcium-binding adaptor molecule 1;
LPS: lipopolysaccharide;
ML: molecular layer;
NOR: novel object recognition;
OEA: oleoylethanolamide;
P: postnatal day;
PB: phosphate buffer;
PBS: phosphate-buffered saline;
PCD: Purkinje cell degeneration;
PCL: Purkinje cell layer;
rhVEGF-B: recombinant human vascular endothelial growth factor B;
RT–qPCR: reverse transcription quantitative polymerase chain reaction;
SCA1: spinocerebellar ataxia type 1;
SD: standard deviation; WT, wild type.

## Declarations

### Ethics approval and consent to participate

All animal experiments were conducted in accordance with European legislation governing the use of laboratory animals, including Directive 2010/63/EU and Recommendation 2007/526/EC, as well as Spanish legislation, including Law 32/2007 and Royal Decree 53/2013. All experimental procedures were approved by the Bioethics Committee of the University of Salamanca, Spain, under reference numbers #00613 and #1258.

### Consent for publication

Not applicable.

### Availability of data and materials

The datasets used and/or analysed during the current study are available upon reasonable request to the corresponding authors.

## Funding

This work was supported by the Spanish Ministry of Economy and Competitiveness (MINECO; SAF2016-79668-R), the Spanish Ministry of Science and Innovation (MICINN; PID2019-106943RB-I00, PID2022-140456NB-I00 and Research Network RED2022-134081-T), and the Regional Government of Castile and León (SA129P20 and SA112P24). D.P.-B. was supported by a predoctoral contract from the University of Salamanca, co-funded by Banco Santander.

### Authors’ contributions

D.P.-B. designed the study, conducted animal experiments and carried out histological and molecular analyses. He performed the statistical analyses, interpreted the results, prepared the figures and wrote the original draft.

J.P.-A. performed CD68 and GFAP immunostaining and the corresponding histological analyses.

J.V. developed Fiji macros for selected image analyses.

E.W. and D.D. contributed to the conception and design of the study, the development of its methodology and the interpretation of the results. They supervised and coordinated the project, helped secure funding and resources, and reviewed and edited the manuscript.

All authors read and approved the final manuscript.

## Acknowledgements

The authors thank María Sánchez for technical support.

