## Supplementary Figure S1 for "Partial microglial depletion attenuates Purkinje cell degeneration and reshapes both glial and inflammatory responses in a mouse model of selective neurodegeneration"

*Supplementary file 1*

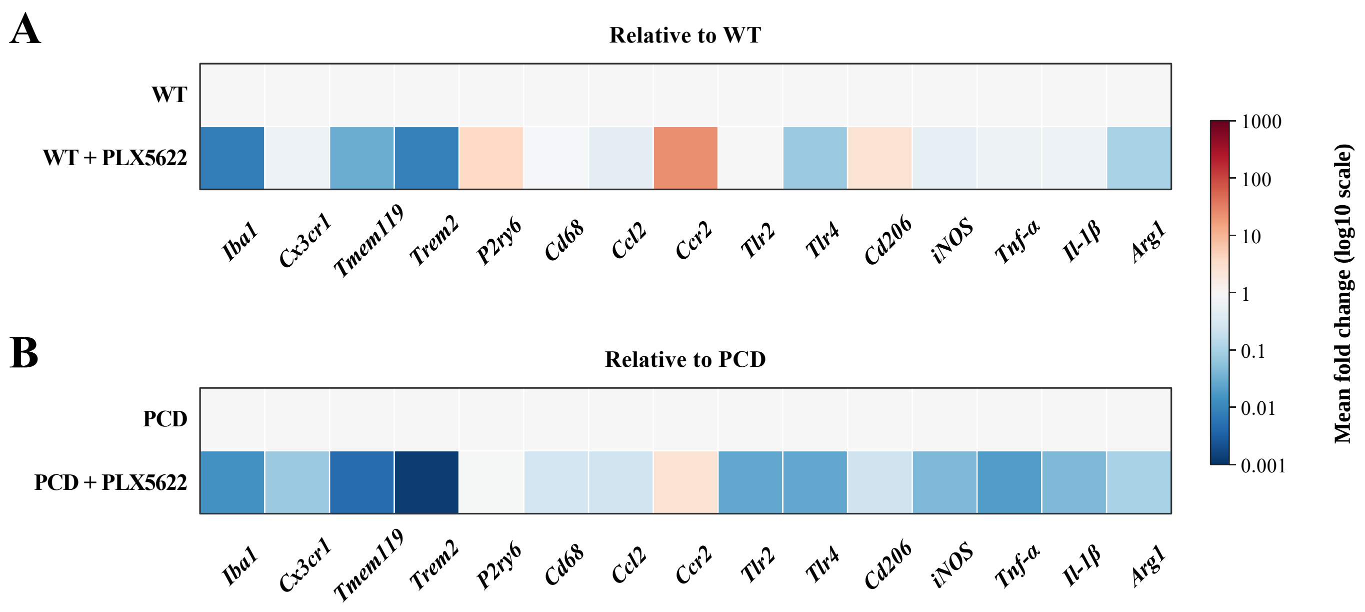


**Supplementary Figure S1. Genotype-matched representation of the cerebellar transcriptional response.**

Mean RT–qPCR fold changes at P25 are shown relative to untreated WT (A) or PCD (B), on a shared logarithmic scale (0.001–1000). Data and statistical comparisons: Figure 8; n = 4 independent animals per group. WT, wild type; PCD, Purkinje cell degeneration.
